# Drug interaction mapping with proximity dependent enzyme recruiting chimeras

**DOI:** 10.1101/2022.09.26.509259

**Authors:** John D Venable, Ajay A Vashisht, Shima Rayatpisheh, James P Lajiness, Dean P Phillips, Ansgar Brock

## Abstract

Proximity dependent labeling using engineered enzymes has been used extensively to identify protein-protein interactions, and map protein complexes *in-vitro* and *in-vivo*. Here, we extend the use of engineered promiscuous biotin ligases to the identification of small molecule protein targets. Chimeric bi-functional chemical probes (“recruiters”) are used to effectively recruit tagged biotin ligases for proximity dependent labeling of target and target interactors. The broad applicability of this approach is demonstrated with probes developed from a multi-kinase inhibitor, a bromodomain targeting moiety, and an FKBP targeting molecule. While complementary to traditional chemo-proteomic strategies such as photo-affinity labeling (PAL), and activity-based protein profiling (ABPP), this approach is a useful addition to the target ID toolbox with opportunities for tunability based on the inherent labeling efficiencies of different engineered enzymes and control over the enzyme cellular localization.

## Introduction

It is difficult to advance small molecule compounds to the clinic without understanding their molecular targets and mechanism of action (MoA). Knowledge about a compound’s molecular target is critical for drug discovery programs to improve selectivity and potency, evaluate pre-clinical toxicology, develop robust pharmcokinetics (PK) and -dynamics (PD) and target engagement packages, and prioritize targets in a commercial environment that supports few clinical starts amongst a plurality of targets. Identification of screening hits and initial characterization of structure-activity relationship (SAR) without knowledge of targets and/or MoA has become common again with the resurgence of phenotypic screening, where cellular or whole organism events are read out without necessarily a detailed understanding of involved pathways and their components^*1-4*^. Consequently, strategies to elucidate the molecular targets of compounds (often referred to as target ID) have become increasingly valued.

The systems biology approach using tools such as functional genomics, and genome wide RNA expression profiling coupled with bioinformatics have fundamentally changed our understanding of cellular pathways, and signaling networks, and have been widely used for MoA interrogation^1, 5^. One of the limitations of these strategies, however, is that they routinely identify large numbers of differentially regulated genes, often because of downstream effects of signaling or pathway modulation, resulting in uncertainty in the assignment of efficacy targets and MoA. Therefore, orthogonal molecular interaction focused chemo-proteomic approaches, such as affinity-based pulldowns^*6*^, photo-affinity labeling^*7*^, activity-based protein profiling^*8*^, and more recent methods like µMapping^9^ have been used extensively to tackle these challenges by providing information on physical protein-drug interactions. While these approaches are invaluable tools for exploration of the target ID landscape, additional orthogonal tools are needed to tackle the unpredictable and fastidious nature of target ID. For example, most covalent probes utilized in ABPP workflows target active site serine and cysteine residues and can therefore only be applied to a subset of targets. Similarly, PAL with minimalist diazirine linkers suffers from low conjugation efficiency as most probe molecules react with solvent, severely limiting sensitivity^*9*^. With these shortcomings in mind, an approach that provides generic targeting and simultaneously allowing for signal amplification based on proximity-dependent enzymatic labeling was devised.

Proximity-dependent labeling (PL) using engineered enzymes in living organisms has emerged as a powerful method to dissect protein-protein interactions and cellular localization^*10*^. PL approaches work by covalently attaching biotin to proteins in the immediate environment of the reagent generating enzyme based on spatial and/or reagent life-time constraints. Enzymes used for this purpose are typically engineered or wild-type versions of biotin ligases or peroxidases, even so an artificial nucleic acid-based mimic of horseradish peroxidase has also been described in the literature^*11*^. Biotin ligase-based enzymes are engineered BirA variants that tag neighboring lysine residues with biotin and include BioID^*12*^, BioID2^*13*^, BASU^*14*^, TurboID^*15*^, Split-BioID^*16*^, and Split-TurboID^*17*^. These enzymes cover a range of kinetics which afford flexibility in optimizing signal to noise and temporal resolution by adjusting labeling times.

Biotin ligases have the advantage of offering consistent activities throughout the cellular environment. In contrast, peroxidase-based enzymes including APEX^*18*^, APEX2^*19*^, and HRP^*20*^ utilize peroxide and biotin-phenol to generate phenoxyl radicals that covalently modify electron-rich residues such as tyrosine. Importantly, the labeling time with peroxidase enzymes (∼1min) is shorter than that of most biotin ligases which provides improved temporal resolution. In practice, PL enzymes are typically used as protein-fusions with “bait” proteins and the resulting biotinylated interacting proteins are subsequently enriched and identified using affinity capture and mass spectrometry.

In this work, we expand PL from protein-fusion based to low molecular weight (LMW) guided for direct identification of protein targets of lead compounds (Fig. 1a). Our strategy, ligand mediated proximity labeling (LMPL), is conceptually similar to the PROTAC approach, where a bi-functional molecule is used to drive target protein / E3 ligase interaction and subsequent ubiquitination of lysine on interacting proteins. However, in this work, we drive target protein of interest (POI) / PL enzyme interaction utilizing proximity enzyme recruiting chimeras (PERCs) resulting in proximity-dependent biotin labeling of interacting proteins. The chimeric recruiter molecules contain (1) a PL enzyme targeting functionality, (2) linker, and (3) POI targeting moiety. Importantly, the desired PL enzyme must be expressed in the cell line of interest. This approach can provide a history of recruiter associations over time (capturing transient or weak interactions) or provide a snapshot of interactions over a brief time depending on the kinetics of the PL enzyme employed. A similar strategy based on direct covalent tagging utilizing the Nedd8 enzyme shows small molecule directed labeling with a fusion protein in lysates is indeed feasible^*21*^.

**Figure 1.**
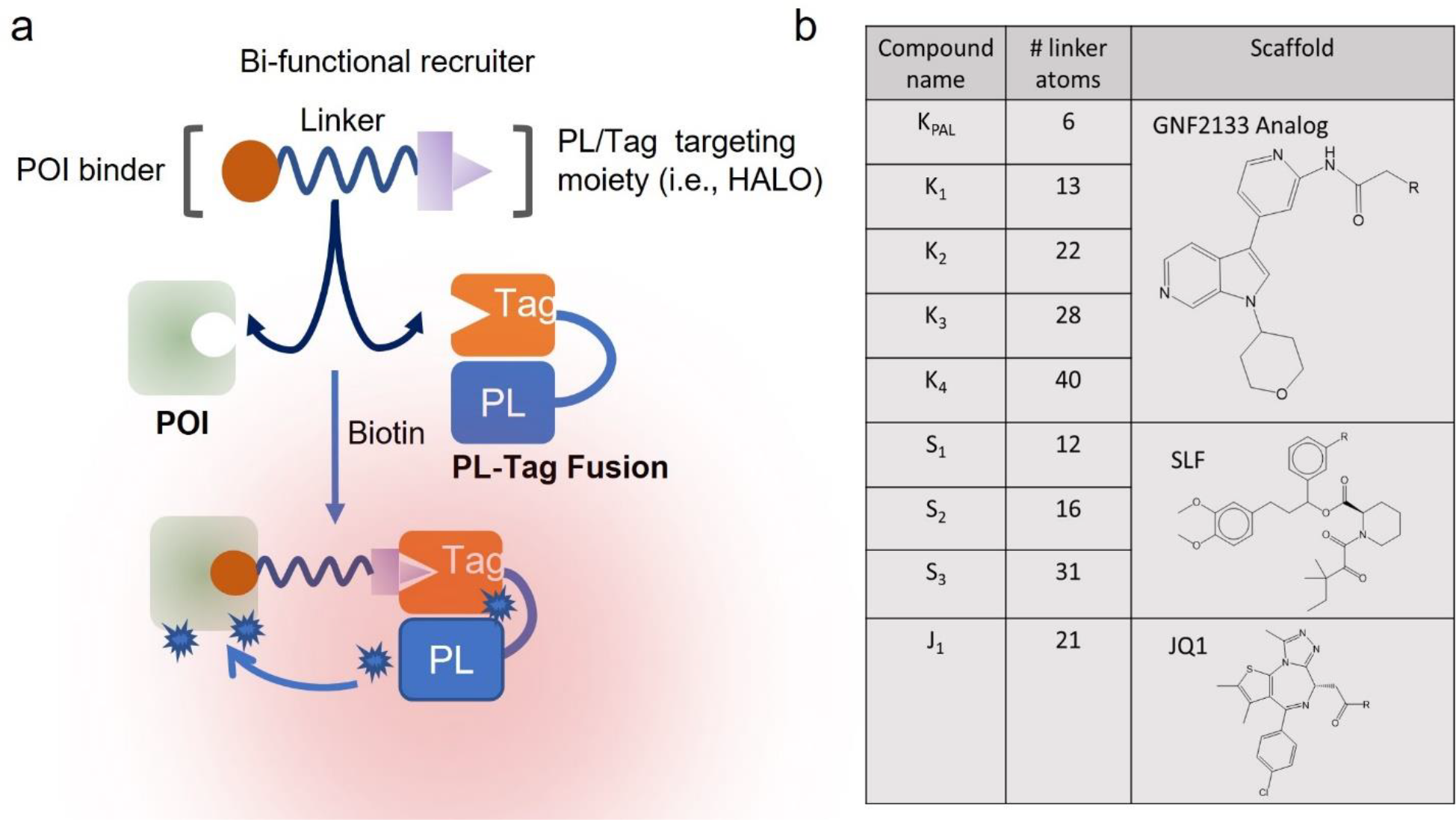
(a) Schematic of the proximity enzyme recruiter concept that utilizing a bi-functional ligand to bring the proximity labeling enzyme (PL) into proximity of target proteins (POI). The example shows a Halo fusion, but other fusion tags including SNAP, CLIP, or FKBP12 (as used in the dTAG strategy) could serve as recruitment handles. Additionally, PL enzymes could be targeted directly with appropriate chemical matter. (b) Chemical probes / recruiters derived from a multi-kinase inhibitor (GNF2133), the FKBP targeting moiety SLF, and the bromodomain targeting compound JQ1. See Supporting Information Table S1 for a full description of the linker-recruitment groups R.

Here, we evaluate conditions for the successful implementation of LMPL and provide several proof-of-concept examples using recruiters that were derived from small molecules with validated molecular targets. For expediency, we chose to generate BioID2, as well as TurboID Halo^22^ & SNAP^23^ tag fusions and utilize chimeric molecules with Halo and SNAP ligands as recruitment handles, although one could screen PL enzymes to identify direct ligand binding sites if a fully reversible recruiter binding mode was desired. Importantly, competition with parent molecules and quantitative proteomics were used to elucidate specifically interacting target proteins from background labeled proteins just as with PAL, and ABPP workflows.

## Results & Discussion

### Design of PL Fusion Constructs and Chimeric Recruiter Compounds

For proof-of-concept studies, the BioID2^13^ & TurboID^15^ enzymes were cloned with Halo and SNAP tags in a HEK293 Flip-in expression system (Supporting Information Fig S1a) and stable cell lines expressing all 4 constructs were generated. The enzymatic activities of the stably expressing constructs were checked by addition of biotin (12.5uM) to cells in 6 well-plates and subsequent immunoblot with streptavidin-HRP (Supporting Information Fig S1b). The results showed self-biotinylation with both the BioID2 and TurboID constructs as well as labeling of many background proteins. The non-specific background labeling was significantly higher for TurboID constructs in accordance with previous studies^*15*^. The activity in freeze-thaw generated lysates was also verified across a biotin labeling time-course with increasing levels of self-biotinylation evident for both constructs across the time-course. The TurboID construct again appeared to be significantly more active than the BioID2 enzyme as evidenced by the increased self, and non-specific background biotinylation at longer time-points.

We next generated a series of recruiter molecules based on several literature reported protein small molecule interaction examples to evaluate the LMPL approach (Fig 1b and Supporting Information Table S1). These model systems included probes based on an analog of the kinase inhibitor GNF2133^*24*^, the FKBP targeting molecule SLF^*25*^, and the BRD 2/3/4 targeting moiety JQ1^*26*^.

The HaloPROTAC system provided a valuable starting point for the development of recruiter molecules^27^ and offered several advantages including speed, the ability to easily monitor Halo tag engagement, and the wealth of information already published. Recent studies have shown the potential for streamlining probe designs so that both PAL/ABPP and recruiter studies could be executed utilizing the same chemical probes that incorporate a chloroalkane^*28*^. Based on the published HaloPROTAC3 optimal linker lengths^*27*^, we synthesized compound K_1_ as well as several extended linker versions K_2_-K_4_ terminating with the Halo tag reactive chloroalkane. SNAP liganded versions K_5_-K_7_ (Supporting Information Table S1) were synthesized for comparison and used in cells expressing PL-SNAP tagged constructs. We also synthesized several chloroalkane terminating recruiter molecules S_1_-S_3_ with varying linker lengths based on the SLF (synthetic ligand for FKBP) moiety, and lastly a recruiter J_1_ based on the well-established BRD2/3/4 targeting ligand JQ1.

### Kinase Inhibitor Model System

Our initial evaluations focused on optimization of labeling conditions, as well as the role of linker length and permeability in the kinase inhibitor system. We therefore decided to use PAL to focus our efforts on the key molecular targets of a GNF2133 analog utilizing a corresponding PAL probe with a minimal diazirine linker and alkyne handle K_PAL_. The PAL results confirmed enrichment of several kinases including the putative target DYRK1A, as well as GSK3A/GSK3B, and several CDK family members (Supporting Information Fig S2a). Additionally, GSK3A/B were among the top hits utilizing K_PAL_ in a chemo-proteomic approach for kinase selectivity profiling (Supporting Information Fig S2b). Therefore, we decided to use a targeted experiment focusing on GSK3A/B to evaluate the labeling efficiencies of the recruiter compounds. To that end, we treated cells that were stably expressing TurboID-Halo, and TurboID-SNAP constructs with 1µM K_1_-K_4_, and K_5_-K_7_ respectively followed by streptavidin bead capture and immunoblotting with anti-GSK3A/B antibody (Supporting Information Fig S2c). The resulting band intensities for GSK3A/B were strongly correlated with linker length suggesting the trimeric complex (PL-Tag / recruiter / GSK3A/B) is inefficiently formed with linkers that are too short. Additionally, the GSK3A/B band intensities were significantly greater with the TurboID-Halo construct than with the TurboID-SNAP version. It was unclear whether the measured GSK3A/B labeling differences were attributable to the protein tags themselves (for example tag geometry) or to physical properties of the probes, so we decided to focus our remaining efforts on Halo tagged constructs and corresponding recruiter probes as they provided the most sensitivity in the kinase inhibitor model system.

To look more closely into the role of linker length on trimeric complex formation, we used an HTRF assay with purified biotinylated DYRK1A kinase domain, and commercially available Halo-GST protein in conjunction with recruiters K_1_-K_4_. The results agreed with the trends seen in the GSK3A/B immunoblotting, and showed trimeric complex formation was strongly impacted by linker length (Fig 2a). In fact, linkers ≥ 28 atoms seemed to be able to saturate complex formation. With concerns about these long linker molecules in mind, we next looked at the permeabilities of the recruiters in our cellular systems.

**Figure 2.**
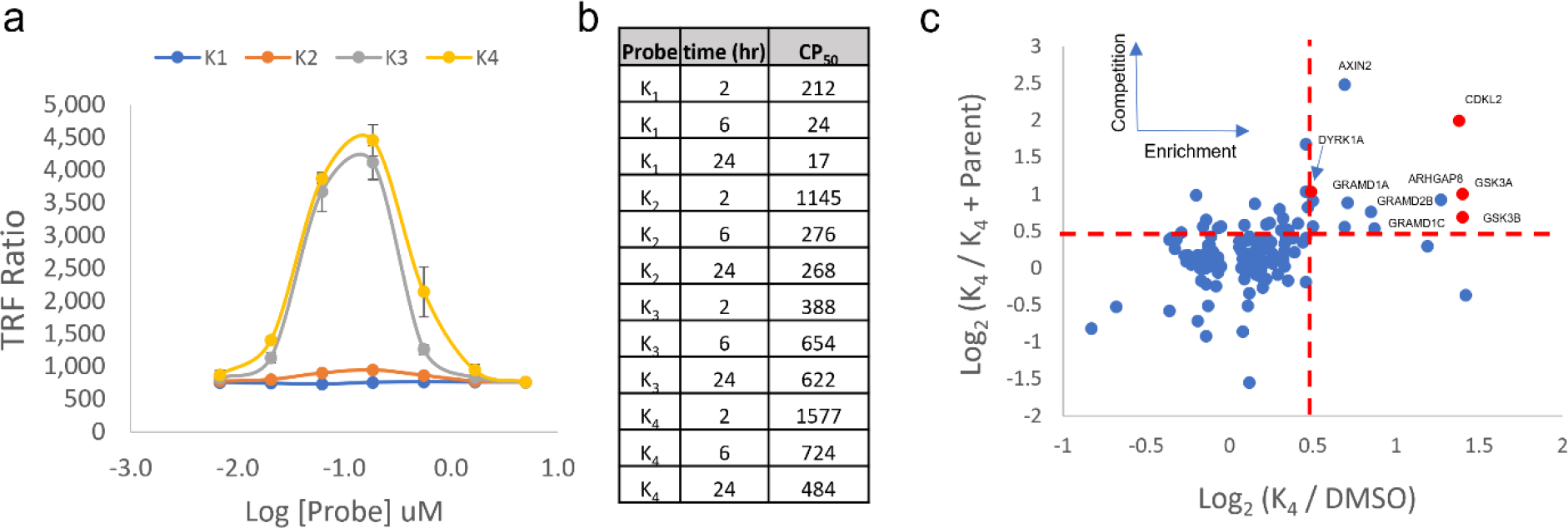
(a) HTRF assay showing impact of linker length on trimeric complex formation with biotinylated Dyrk1a kinase domain, commercially available Halo-GST protein, and bi-functional recruiter probes. (b) Measured CP50 (µM) using the chloroalkane penetration assay showing the impact of linker length on compound permeability. (c) 2D scatter plot showing protein enrichment vs. competition utilizing probe K_4_ in TurboID-Halo expressing cells and at a 2hr labeling time-point (only points with p-value <0.05 shown, kinases highlighted in red).

We evaluated the permeability of K_1_-K_4_ using the chloroalkane penetration assay^*29*^, at three time-points (2, 6, and 24hr) and found all three compounds were able to penetrate cells stably expressing the TurboID-Halo construct and covalently react with the Halo tag. The CP_50_’s (concentration at which 50% of the maximal cell penetration was observed, Fig 2b) correlated with linker length and the longer linker recruiters required significantly higher concentrations and/or longer labeling times to effectively label the Halo tag. In our subsequent LMPL studies, we therefore chose to use recruiter concentrations at least 1.5-fold greater than the CP_50_, and minimum labeling time-points of 2 hr.

### Kinase Target Deconvolution

In our initial studies of the LMPL strategy, we compared the enrichment of kinases utilizing the recruiter K_4_ in cell lines stably expressing BioID2-Halo and TurboID-Halo constructs (Supporting Information Fig. S3a-b) at commonly used time-points, and biotin concentrations for these enzymes. We saw several known kinase targets of the GNF2133 warhead were enriched in the TurboID system, but not in the BioID2 system. We surmised that the increased activity of the TurboID enzyme was responsible for the difference and proceeded to run competition studies utilizing K_PAL_ to directly compete with K_4_ in TurboID-Halo expressing cells at multiple labeling time-points (Fig. 2c, Supporting Information Fig. S4).

In line with previous studies using TurboID^*15*^, we saw that the labeling time-point had a large effect on the number of proteins that were enriched (Log2 [recruiter / DMSO] >0.5, p-value < 0.05) in the pulldown experiments, with 13, 22, and 626 proteins quantified after 2, 6, and 24h respectively (Supporting Information Fig. S3b). The differences are likely attributable to increased background labeling, and care should be taken to work at the shortest time-point where sufficient signal is acquired (2-6hr in these studies). Several known kinase targets of this warhead including DYRK1A, GSK3A/B, CSNK1D, CDK’s and CDKL2 passed our hit picking criteria (Log2 [recruiter / DMSO] > 0.5, Log2 [recruiter / recruiter + PAL] > 0.5) validating LMPL as a viable strategy for Target ID. The enrichment of kinases in the LMPL experiments (Log_2_ [recruiter / DMSO]) were lower than that obtained in the corresponding PAL experiment (Supporting Information Fig S2a), however the enrichment of background proteins was also significantly lower resulting in strong signal to noise, and the potential to use lower S/N thresholds for “hit” prioritization.

Interestingly, several known cytosolic kinase targets of the warhead were enriched in the LMPL experiment; however, known nuclear kinases targets were significantly underrepresented which might be due to primarily cytosolic localization of the PL enzyme. We therefore envision future experiments to interrogate the role of PL construct localization in LMPL and ascertain if localization can be used to preferentially enrich targets based on their sub-cellular localization.

### SLF Target Deconvolution

As another proof-of-concept example, we utilized probes S_1_-S_3_ based on the SLF moiety which is a known inhibitor of FKBP family members including FKBP5 (affinity ∼3.1uM), and FKBP1A (affinity ∼2.6uM). Two replicate pulldowns were performed to evaluate enrichment and a competition study with a 10X excess of the parent molecule, SLF, was also performed.

Unfortunately, FKBP1A was not detected in either experiment, however we saw strong enrichment of FKBP5 with recruiter S_2_ in both replicate pulldowns (Fig 3a-b). Interestingly, lower enrichment of FKBP5 (Fig 3c) was obtained with probes S_1_ and S_3_ suggesting the optimal linker length is ∼16 atoms for enrichment in this context. Importantly, competition with a 10-fold excess of SLF decreased the enrichment of FKBP5 significantly (Fig 3d) suggesting FKBP5 biotinylation by TurboID-Halo is specific and driven by the recruiter molecule.

**Figure 3.**
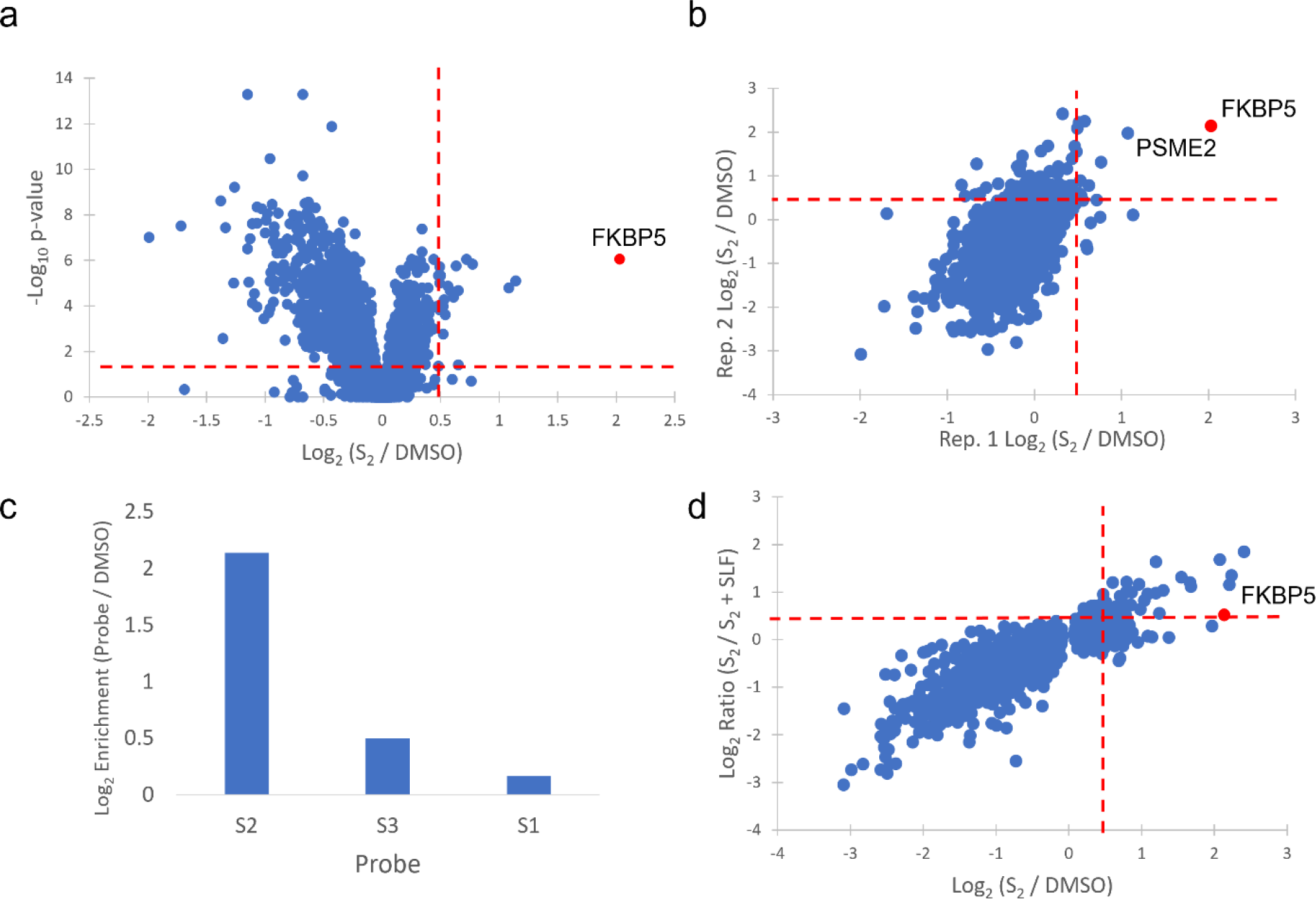
(a) Volcano plot showing replicate #1 of a pulldown utilizing S_2_ (2.5µM) in TurboID-Halo expressing cells at a 3hr time-point (b) Correlation plot showing enrichment in replicate pulldown experiments (c) Enrichment of FKBP5 with different recruiter probes (d) 2D scatter plot showing protein enrichment vs. competition utilizing probe S_2_ in TurboID-Halo expressing cells and at a 3hr labeling time-point.

### JQ1 Target Deconvolution

Lastly, we used the recruiter J_1_ based on the validated bromodomain inhibitor (+)-JQ1, which has been shown to potently inhibit the BET family of bromodomain proteins (BRD 2/3/4). We validated target engagement in cells by treating TurboID-Halo expressing stable cells with 12.5µM biotin, and 1µM J_1_ for 3hr. After lysis and affinity capture of biotinylated proteins, we found that BRD 2/3/4 were all significantly enriched (Log_2_ [recruiter / DMSO] > 0.5, Fig 4a).

**Figure 4.**
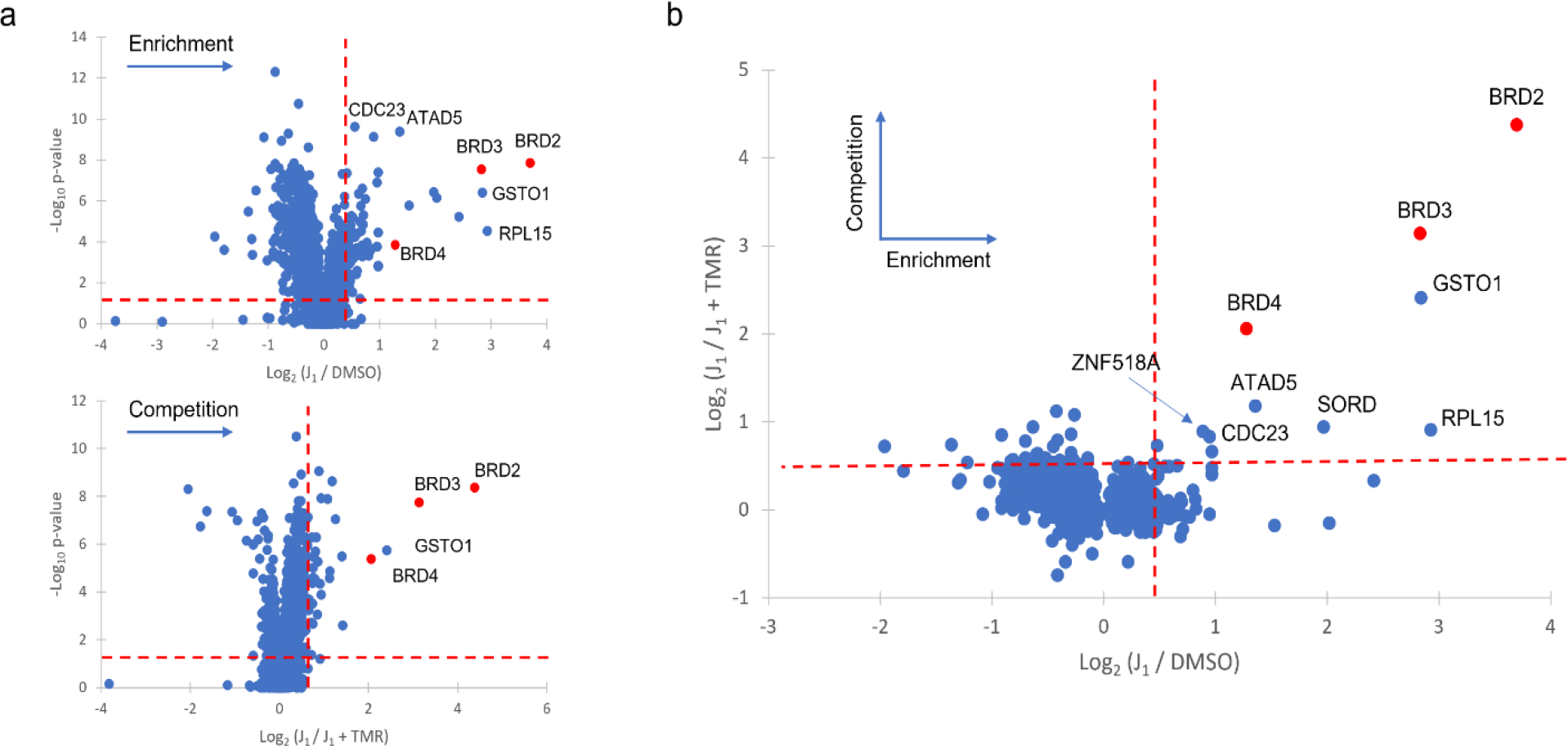
(a) Volcano plots showing enrichment and competition in pulldowns utilizing J_1_ in TurboID-Halo expressing cells and at a 3hr time-point (b) 2D scatter plot utilizing probe J_1_ in TurboID-Halo expressing cells and a 3hr labeling time-point (BRDs highlighted in red).

Importantly, the enrichment was blocked by pre-treatment of the cells with the commercially available TMR-Halo ligand, which blocks the HaloTag binding site, and shows that labeling is specific to the Halo tagged proximity enzyme construct (Fig 4b). In agreement with expectations, several known bromodomain interacting proteins including ATAD5^30, 31^, ZNF518A^*32*^, and CDC23^32^ passed hit-picking criteria (Log_2_ [recruiter / DMSO] > 0.5, Log_2_ [recruiter / recruiter + parent] > 0.5) suggesting interacting proteins and complex members are biotinylated during the labeling procedure. LMPL therefore provides an additional tool to study protein complex stabilization or disruption upon treatment with small molecule probes and could be particularly valuable for targeted protein stability and/or targeted protein degradation studies.

## Conclusions

In this work, we describe a strategy for interrogating protein-small molecule interactions using low molecular weight guided recruitment of engineered biotin ligases. We show that this strategy facilitates identification of protein targets as well as protein complex members in proximity to targeted proteins. The proof-of-concept studies used the HaloTag system along with well-studied engineered biotin ligases for expediency. However, we view the tunability of this approach as an asset which affords exciting new opportunities including the potential mapping of recruiter targets and target interacting proteins in-vivo using genetic knock-in expression of PL enzymes^*33*^. Future efforts should evaluate the potential benefits of other engineered enzymes, the role of enzyme sub-cellular localization, control of level of expressed fusion proteins, and the merit of a reversible binding system that could promote enhanced catalytic activity among other things. In conclusion, we anticipate the LMPL strategy will complement existing chemo-proteomic methods while also providing a powerful and tunable approach for dissecting protein-small molecule interactions.

## Materials / Methods

### Cloning and generation of stable cell lines

Primer and sequence information of the various constructs that were designed for this study are shown in Supporting Information Table S2. Gene blocks for SNAP, Halo, BioID2 and TurboID were obtained from IDT. Each fragment was PCR amplified with primers shown in Table S2 and cloned into mammalian expression vector pcDNA5FRT/TO using Gibson assembly that led to generation of the following fusion partners: BioID2-SNAP, TurboID-SNAP, BioID2-Halo and TurboID-Halo at HindIII and XhoI sites. Each construct was verified by restriction digestion analysis and sequencing. Plasmid constructs were used to obtain stable cell lines in Flp-InTM T-RExTM-293 cells (Thermo Scientific). The cells were cultured and maintained as described by the manufacturer. Flp-InTM T-RExTM-293 cells stably expressing BioID2-Halo, TurboID-Halo, BioID2-SNAP, and TurboID-SNAP constructs were generated using the Flp-In system (Invitrogen) as per manufacturer’s instructions. The expression of individual constructs in stable Flp-InTM T-RExTM-293 cells was driven by adding doxycycline overnight to a final concentration of 500 ng/mL.

### Photo-affinity labeling

15cm plates of HEK293 cells were grown to confluence, and cells were harvested by scraping. Cells were washed 2X in ice cold PBS and lysed in ice cold Lysis Buffer (20mM Hepes, pH 7.5, 1% NP40, 1X HALT protease inhibitor cocktail). Pierce universal nuclease (Thermo Scientific) was added, and lysate was incubated on ice for 60 min with occasional mixing. Lysate was centrifuges at 14,000 g for 15min at 4°C to pellet cell debris. The supernatant was removed and aliquoted into 2mL Eppendorf tubes for further processing. A pierce 660nm protein assay was performed to determine protein concentration.

5mg of total lysate was used for each treatment condition. Pre-clearing of lysate to remove endogenously biotinylated proteins was performed using 250ul of high-capacity streptavidin agarose beads (Pierce) packed into 2mL Pierce centrifuge columns. Lysate was incubated with beads for 1hr at 4°C while rotating, and then centrifuged for 3min at 300g to elute the pre-cleared lysate. Columns were washed 1X with 0.5mL Lysis Buffer and combined with previously eluted lysate.

DMSO was added to the control lysate sample, and PAL probe K_PAL_ was added to the treatment sample at a final concentration of 500nM and incubated for 2hr at 4C while rotating. Lysates were transferred to 6 well plates and UV crosslinked using a Spectrolink 1500 at 365nM with an integrated plate cooler held at 4°C. Crosslinked lysates were subjected to copper catalyzed click reaction using pre-mixed CuSO_4_ and Tris(3-hydroxypropyltriazolylmethyl)amine (Sigma) at 2.5µM and 1.25 µM respectively, 1µM picolyl-biotin-azide (Click Chemistry Tools), and 2.5µM sodium ascorbate (Sigma) for 1hr at room temperature while rotating. Samples were then incubated with 250uL pre-washed streptavidin mag-sepharose (Cytiva) beads for 1 hr. at room temperature while rotating. Beads were washed, digested, TMT labeled, and fractionated as described in the Supporting Information.

### Ligand Mediated Proximity Labeling

#### Kinase targeting recruiter cell treatments

BioID-Halo: 15cm plates of stably expressing BioID2-Halo cells were plated and allowed to attach for 2hr. 1ug/mL Doxycycline was added to all plates and cells were incubated 24 hr. DMSO or recruiter compounds (2.5µM) were then added followed immediately by 25 µM biotin. After 24hr, the plates were harvested by scraping.

TurboID-Halo: 15cm plates of stably expressing TurboID-Halo cells were plated and allowed to attach to plates for 2hr. 1ug/mL Doxycycline was added to all plates and cells were incubated for 24hr. Recruiter compounds (2.5µM) were then added followed immediately by 12.5 µM biotin. For competition, K_PAL_ was added at a 10-fold excess 30min prior to the recruiter compounds. Cells were incubated for labeling time-points of 2, 6, and 24hr and then harvested by scraping.

#### SLF based recruiter cell treatments

15cm plates of stably expressing TurboID-Halo cells were plated and allowed to attach for 2hr. 1ug/mL Doxycycline was added to all plates and cells were incubated for 24hr. DMSO or recruiter compounds S_1_-S_3,_ (2.5µM) were then added followed immediately by 12.5 µM biotin. For competition, the parent molecule SLF (25 µM) was added 30min prior to the recruiter compounds. Cells were incubated for a labeling time-point of 3hr and plates were harvested by scraping.

#### JQ1 based recruiter cell treatments

15cm plates of stably expressing TurboID-Halo cells were plated and allowed to attach for 2hr. 1ug/mL Doxycycline was added to all plates and cells were incubated for 24hr. Recruiter compound J_1_ (1µM) was then added followed immediately by 12.5 µM biotin. For competition, the TMR-Halo ligand was added 60min prior to recruiter compound at 5µM. Cells were incubated for a labeling time-point of 3hr, and plates were harvested by scraping.

#### Biotinylated protein pulldowns

Harvested cells were washed 2X in ice cold PBS, pelleted by centrifugation, and lysed in 1.5 mL ice cold Lysis Buffer (20mM Hepes, pH 7.5, 1% NP40, 1X HALT protease inhibitor cocktail).

Pierce universal nuclease (Thermo Scientific) was added, and lysate was incubated on ice for 60 min with occasional mixing. Lysate was centrifuges at 14k g for 15min at 4°C to pellet cell debris. The supernatant was removed and aliquoted into 2mL Eppendorf tubes for further processing. A 20ul aliquot was removed for 660nm protein assay for total protein normalization. Streptavidin mag-sepharose (Cytiva) beads were washed 2X with Lysis Buffer and 250uL beads were mixed with lysate from each condition. Samples were incubated at room temperature for 1hr while rotating. Beads were washed, digested, TMT labeled, and fractionated as described in the Supporting Information.

## Acknowledgements

The authors would like to thank Markus Schirle for his help editing the manuscript.

