## Supporting Information for "Drug interaction mapping with proximity dependent enzyme recruiting chimeras"

**Supporting Figures**

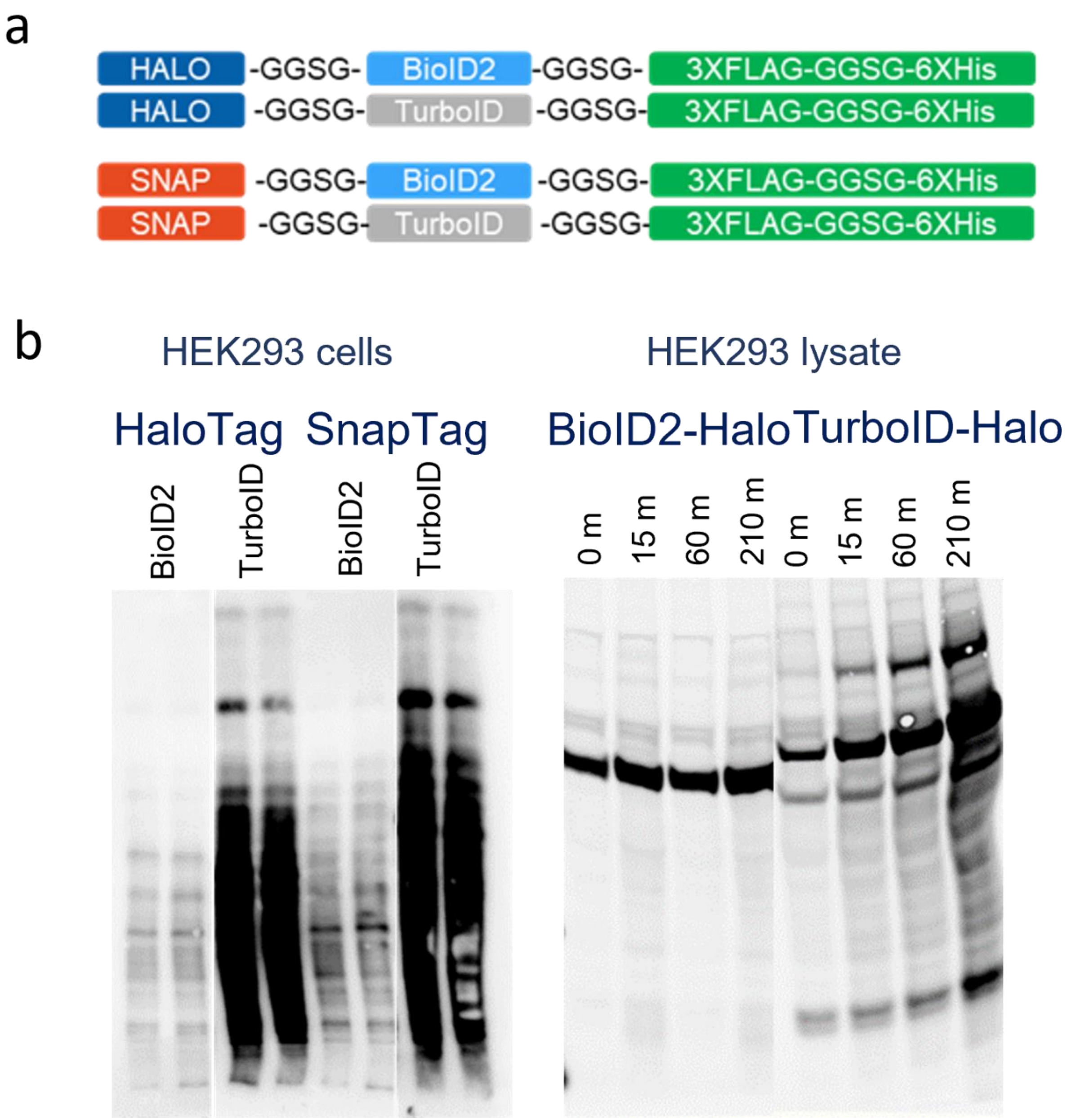

**Figure S1.** (a) Schematic of constructs cloned with Halo and SNAP tags in a HEK293 Flip-in expression system (b) enzymatic activities of the stably expressing constructs in cells and after freeze/thaw lysis checked by streptavidin western.

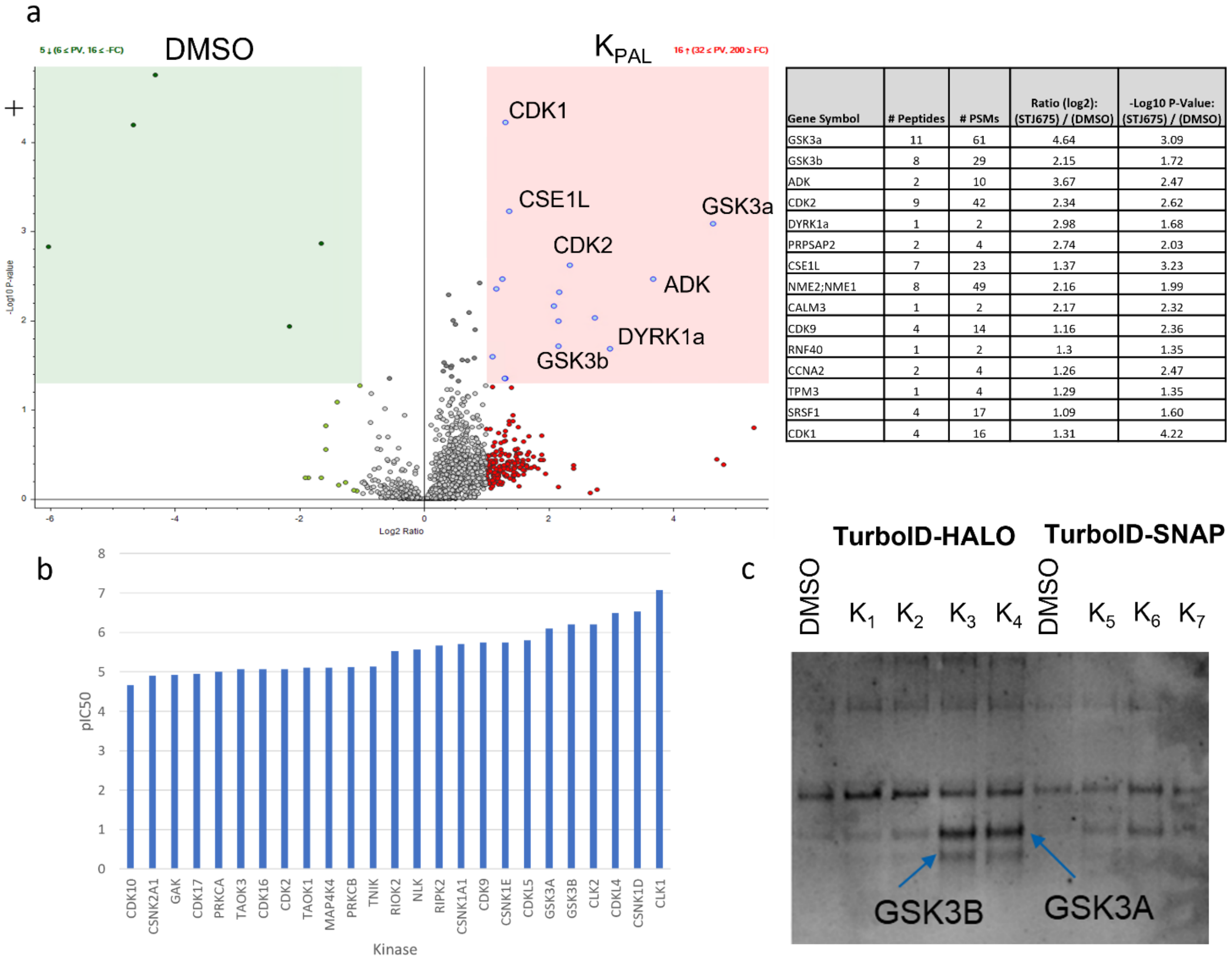

| Gene Symbol | # Peptides | # PSMs | Ratio (log2): (STJ675) / (DMSO) | -Log10 P-Value: (STJ675) / (DMSO) |
|---|---|---|---|---|
| GSK3a | 11 | 61 | 4.64 | 3.09 |
| GSK3b | 8 | 29 | 2.15 | 1.72 |
| ADK | 2 | 10 | 3.67 | 2.47 |
| CDK2 | 9 | 42 | 2.34 | 2.62 |
| DYRK1a | 1 | 2 | 2.98 | 1.68 |
| PRPSAP2 | 2 | 4 | 2.74 | 2.03 |
| CSE1L | 7 | 23 | 1.37 | 3.23 |
| NME2;NME1 | 8 | 49 | 2.16 | 1.99 |
| CALM3 | 1 | 2 | 2.17 | 2.32 |
| CDK9 | 4 | 14 | 1.16 | 2.36 |
| RNF40 | 1 | 2 | 1.3 | 1.35 |
| CCNA2 | 2 | 4 | 1.26 | 2.47 |
| TPM3 | 1 | 4 | 1.29 | 1.35 |
| SRSF1 | 4 | 17 | 1.09 | 1.60 |
| CDK1 | 4 | 16 | 1.31 | 4.22 |

**Figure S2.** (a) Volcano plot from PAL study with GNF2133 analog probe $K_{PAL}$ shows enrichment of kinases including DYRK1a, CDK's, and GSK3a/b (b) Chemo-proteomic kinase selectivity profiling of $K_{PAL}$ in HEK293 lysate (c) Anti-GSK3a/b western after streptavidin capture with probes containing Halo and SNAP ligands in both TurboID-Halo and TurboID-SNAP tagged stable cell lines respectively.

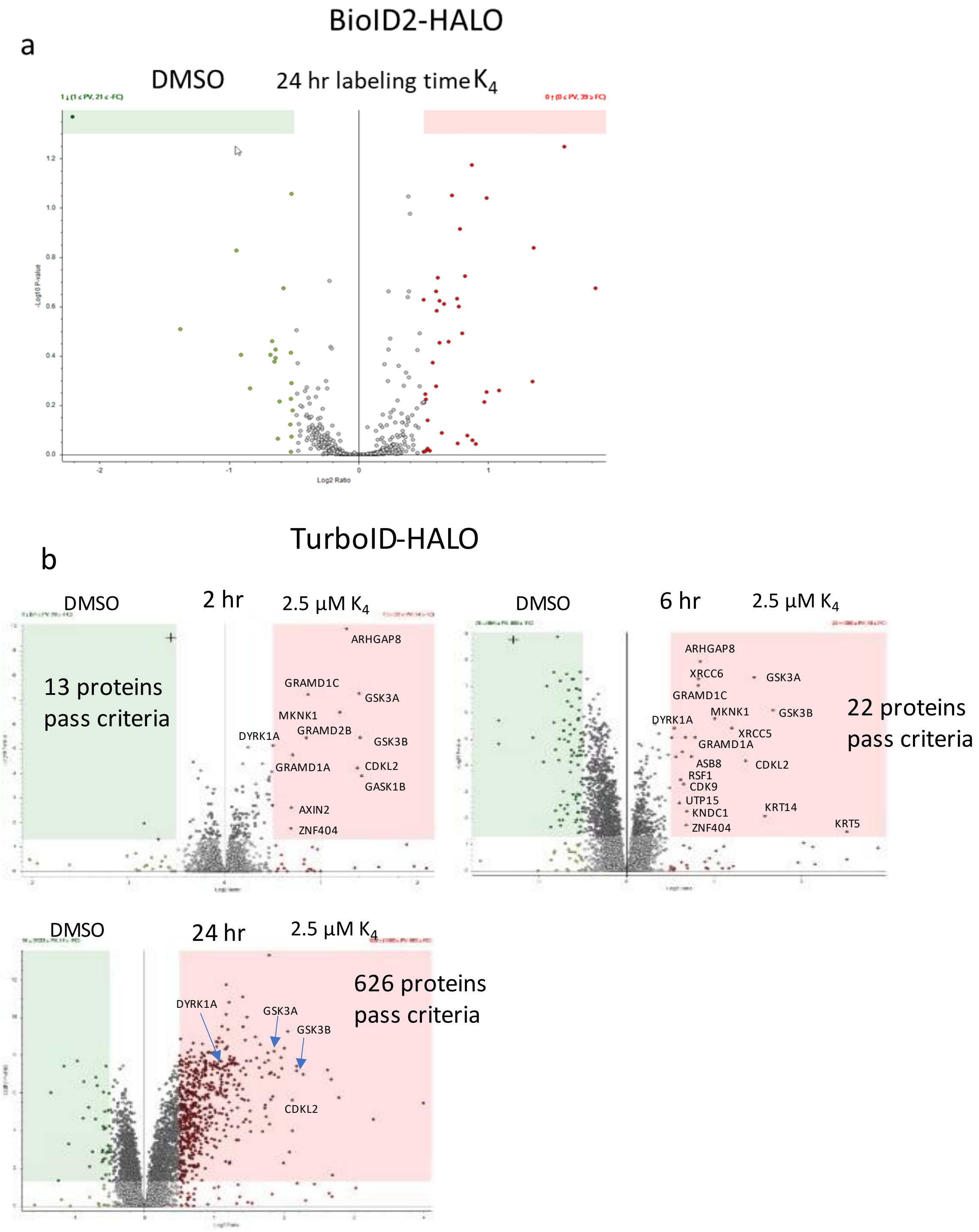

**Figure S3. (a) Volcano plot showing enrichment of proteins utilizing the recruiter $K_4$ in cell lines stably expressing (a) BioID2-Halo and (b) TurboID-Halo constructs at a range of commonly used time-points for these enzymes. Pink and green shading signify highlight proteins that pass filtering criteria ($Log_2$ [$K_4$ / DMSO] > 0.5, and p-value <0.05).**

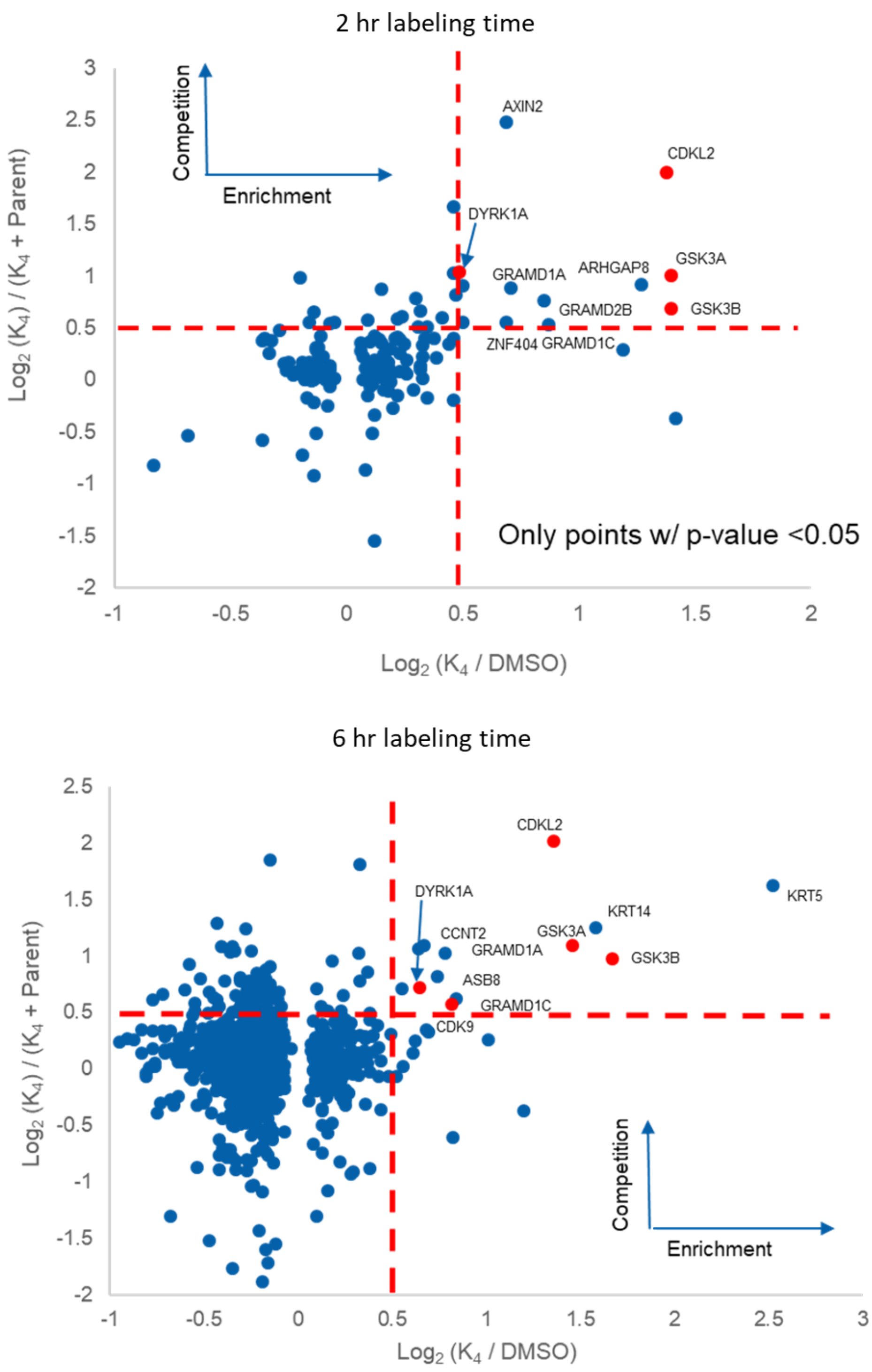

**Figure S4. (a) 2D scatter plot showing protein enrichment vs. competition utilizing probe $K_4$ in TurboID-Halo expressing cells at 2, and 6 hr labeling times.**

**Supporting Table S1: Series of recruiter probes based on literature reported protein small molecule interaction examples.**

| Compound # | # linker atoms | Scaffold | $R_1$ (linker) | $R_2$ (reactive group) |
|---|---|---|---|---|
| $K_{PAL}$ | 6 | GNF2133 Analog 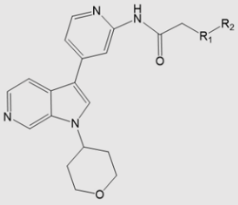 | 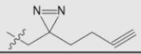 | n.a. |
| $K_1$ | 13 | | 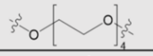 | 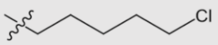 |
| $K_2$ | 22 | | 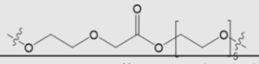 | |
| $K_3$ | 28 | | 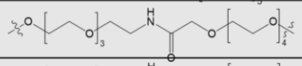 | |
| $K_4$ | 40 | | 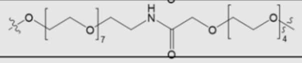 | |
| $K_5$ | 7 | | 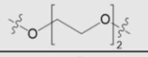 | 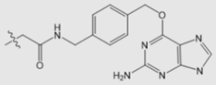 |
| $K_6$ | 22 | | 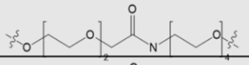 | |
| $K_7$ | 34 | | 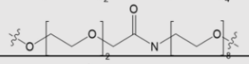 | |
| $S_1$ | 12 | SLF 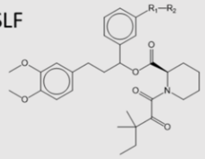 | 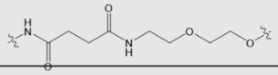 | 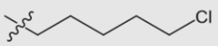 |
| $S_2$ | 16 | | 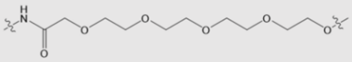 | |
| $S_3$ | 31 | | 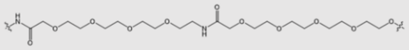 | |
| $J_1$ | 21 | JQ1 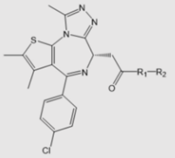 | 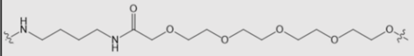 | |

### Supporting Table S2: Primer and sequence information of the various constructs that were designed for this study.

***HALO-BioID2 in pCDNA5FRT/TO: digest vector with HindIII /Xho1***

| Primer name | Sequence | Length |
| --- | --- | --- |
| *F1_5_HALO_BioID_Fwd* | CGGGACCGATCCAGCCTCCGGACTCTAGCGTTTAAACTTAAGCTTATGGCAGAAATCGGTACTGGC | 66-mer |
| *F1_5_ HALO _BioID_Rev* | CAGCCAGATCAGGTTCTTGAAaccagagcctccGCCGGAAATCTCGAGC | 49-mer |
| *F2_5_ HALO _BioID_Fwd* | ATTTCCGGCggaggctctggtTTCAAGAACCTGATCTGGCTGAAGG | 46-mer |
| *F2_5_ HALO _BioID_Rev* | TTAAACGGGCCCTCTAGACTCGAGTCAATGGTGATGGTGATGATGaccagagcctccCTTGTCATCGTCATCCTTGTAATCGATGTC ATGATCTTTATAATCACCGTCATGGTCTTTGTAGTCaccagagcctccGCTTCTTCTCAGGCT | 150-mer |
| ***HALO TurboID in pCDNA5FRT/TO: digest vector with HindIII /Xho1*** | | |
| *F1_5_ HALO _TurboID_Fwd* | CGGGACCGATCCAGCCTCCGGACTCTAGCGTTTAAACTTAAGCTTATGGCAGAAATCGGTACTGGC | 66-mer |
| *F1_5_ HALO _TurboID_Rev* | CAGAGGCACAGTATTGTCTTTaccagagcctccGCCGGAAATCTCGAGCG | 50-mer |
| *F2_5_ HALO _TurboID_Fwd* | ATTTCCGGCggaggctctggtAAAGACAATACTGTGCCTCTGAAGCT | 47-mer |
| *F2_5_ HALO _TurboID_Rev* | GTTTAAACGGGCCCTCTAGACTCGAGTCAATGGTGATGGTGATGATGaccagagcctccCTTGTCATCGTCATCCTTGTAATCGATG TCATGATCTTTATAATCACCGTCATGGTCTTTGTAGTCaccagagcctccCTTTTCGGCAGACC | 151-mer |
| ***SNAP BioID2 in PCDNA5FRT/TO: digest vector with HindIII /Xho1*** | | |
| *F1_5_SNAP_BioID_Fwd* | CGGGACCGATCCAGCCTCCGGACTCTAGCGTTTAAACTTAAGCTTATGGACAAAGACTGCGAAATGAAGC | 70-mer |
| *F1_5_SNAP_BioID_Rev* | CAGCCAGATCAGGTTCTTGAAaccagagcctccGCCACCCAGCCCAGGCTT | 51-mer |
| *F2_5_SNAP_BioID_Fwd* | CCTGGGCTGGGTGGCggaggctctggtTTCAAGAACCTGATCTGGCTGAAGG | 52-mer |
| *F2_5_Halo_BioID_Rev* | TTAAACGGGCCCTCTAGACTCGAGTCAATGGTGATGGTGATGATGaccagagcctccCTTGTCATCGTCATCCTTGTAATCGATGTC ATGATCTTTATAATCACCGTCATGGTCTTTGTAGTCaccagagcctccGCTTCTTCTCAGGCT | 150-mer |
| ***SNAP TurboID pCDNA5FRT/TO: digest vector with HindIII /Xho1*** | | |
| *F1_5_SNAP_TurboID_Fwd* | CGGGACCGATCCAGCCTCCGGACTCTAGCGTTTAAACTTAAGCTTATGGACAAAGACTGCGAAATGAAGC | 70-mer |
| *F1_5_SNAP_Turbo_Rev* | AGAGGCACAGTATTGTCTTTaccagagcctccGCCACCCAGCCCAGGCTT | 50-mer |
| *F2_5_SNAP_Turbo_Fwd* | CCTGGGCTGGGTGGCggaggctctggtAAAGACAATACTGTGCCTCTGAAGCT | 53-mer |
| *F2_5_SNAP_TurboID_Rev* | GTTTAAACGGGCCCTCTAGACTCGAGTCAATGGTGATGGTGATGATGaccagagcctccCTTGTCATCGTCATCCTTGTAATCGATG TCATGATCTTTATAATCACCGTCATGGTCTTTGTAGTCaccagagcctccCTTTTCGGCAGACC | 151-mer |

**Materials and Methods:**

**Construct Sequences**

**SNAP tag** (sequence information was obtained from the NEB website):

ATGGACAAAGACTGCGAAATGAAGCGCACCACCCTGGATAGCCCTCTGGGCAAGCTGGAACTGTCTGG
GTGCGAACAGGGCCTGCACCGTATCATCTTCCTGGGCAAAGGAACATCTGCCGCCGACGCCGTGGAAGT
GCCTGCCCCAGCCGCCGTGCTGGGCGGACCAGAGCCACTGATGCAGGCCACCGCCTGGCTCAACGCCTA
CTTTCACCAGCCTGAGGCCATCGAGGAGTTCCCTGTGCCAGCCCTGCACCACCCAGTGTTCCAGCAGGA
GAGCTTTACCCGCCAGGTGCTGTGGAAACTGCTGAAAGTGGTGAAGTTCGGAGAGGTCATCAGCTACA
GCCACCTGGCCGCCCTGGCCGGCAATCCCGCCGCCACCGCCGCCGTGAAAACCGCCCTGAGCGGAAATC
CCGTGCCCATTCTGATCCCCTGCCACCGGGTGGTGCAGGGCGACCTGGACGTGGGGGGCTACGAGGGC
GGGCTCGCCGTGAAAGAGTGGCTGCTGGCCCACGAGGGCCACAGACTGGGCAAGCCTGGGCTGGGT

**HaloTag** (sequence information was obtained from Promega vector pHTC HaloTag® CMV-neo Vector)

ATGGCAGAAATCGGTACTGGCTTTCCATTCGACCCCCATTATGTGGAAGTCCTGGGCGAGCGCATGCAC
TACGTCGATGTTGGTCCGCGCGATGGCACCCCTGTGCTGTTCCTGCACGGTAACCCGACCTCCTCCTACG
TGTGGCGCAACATCATCCCGCATGTTGCACCGACCCATCGCTGCATTGCTCCAGACCTGATCGGTATGGG
CAAATCCGACAAACCAGACCTGGGTTATTTCTTCGACGACCACGTCCGCTTCATGGATGCCTTCATCGAA
GCCCTGGGTCTGGAAGAGGTCGTCCTGGTCATTCACGACTGGGGCTCCGCTCTGGGTTTCCACTGGGCC
AAGCGCAATCCAGAGCGCGTCAAAGGTATTGCATTTATGGAGTTCATCCGCCCTATCCCGACCTGGGAC
GAATGGCCAGAATTTGCCCGCGAGACCTTCCAGGCCTTCCGCACCACCGACGTCGGCCGCAAGCTGATC
ATCGATCAGAACGTTTTTATCGAGGGTACGCTGCCGATGGGTGTCGTCCGCCCGCTGACTGAAGTCGAG
ATGGACCATTACCGCGAGCCGTTCCTGAATCCTGTTGACCGCGAGCCACTGTGGCGCTTCCCAAACGAG
CTGCCAATCGCCGGTGAGCCAGCGAACATCGTCGCGCTGGTCGAAGAATACATGGACTGGCTGCACCA
GTCCCCTGTCCCGAAGCTGCTGTTCTGGGGCACCCCAGGCGTTCTGATCCCACCGGCCGAAGCCGCTCG
CCTGGCCAAAAGCCTGCCTAACTGCAAGGCTGTGGACATCGGCCCGGGTCTGAATCTGCTGCAAGAAGA
CAACCCGGACCTGATCGGCAGCGAGATCGCGCGCTGGCTGTCGACGCTCGAGATTTCCGGC

**BioID 2**: sequence information was obtained from Kim et al.). This sequence is a humanized A. aeolicus BioID2 with R40G. Gene blocks were synthesized for cloning.

Reference: Kim, Dae In, et al. "An improved smaller biotin ligase for BioID proximity labeling." Molecular biology of the cell 27.8 (2016): 1188-1196.

**TurboID**: sequence information as licensed from Stanford University.

Reference: Branon, Tess C., et al. "Efficient proximity labeling in living cells and organisms with TurboID." Nature biotechnology 36.9 (2018): 880-887.

**Immunoblotting:**

**PL enzyme activity assay**

HEK293 Flp-In Halo-TurboID stable cells and HEK293 Flp-In Halo-BioID stable cells lines each were seeded in 2wells of a six well tissue culture plate. Cells were treated with 1ug/ml final concentration of doxycycline for 24 hours and 12.5uM final concentration of biotin overnight. After harvesting and lysing cells in ice cold Flag lysis buffer (100mM Tris-HCL pH 8.0, 150mM NaCl, 5mM EDTA, 5% Glycerol, 0.1% NP40, halt protease and phosphatase inhibitor) with end over end rotation at 4°C for an hour, lysate tubes were centrifuged at 14,000g for 15 minutes and supernatant transferred to new tubes. The samples were mixed one to one with 2X LDS sample buffer (Pierce #84788) with 2X NuPAGE™ sample reducing agent (ThermoFisher Scientific #NP0009) and boiled at 95°C for 5 minutes.

Samples were separated on a NuPAGE™ 4 to 12%, Bis-Tris, 1.5 mm, Mini Protein Gel, 10-well (ThermoFisher Scientific #NP0335BOX) along with 4ul of PageRuler™ Plus Prestained Protein Ladder, 10 to 250 kDa (ThermoFisher Scientific #26619) for 45 minutes at 200 volts in 1X NuPAGE™ MES SDS Running Buffer (ThermoFisher Scientific #NP0002). Proteins were transferred to a nitrocellulose mini membrane (ThermoFisher Scientific #IB23002) using an iBLOT2 gel transfer device (ThermoFisher Scientific#IB21001). The membrane was blocked in Intercept (PBS) Blocking Buffer (LiCor# 927-70001) for 30 minutes. Streptavidin, Alexa Fluor™ 647 conjugate (Invitrogen #S21374) was added 1:1000 in PBS buffer with 0.2% Tween 20 (Sigma #P9416) to the blot for an hour shacking at room temperature. Blot wash washed four times (each 5 minutes) with PBS and analyzed by BioRad ChemidDoc MP imaging system.

**Anti-GSK3a/b western**

15cm plates of HEK293 Flp-In Halo-TurboID stable cells and 15cm plates of HEK293 Flp-In Snap-TurboID stable cells received doxycycline treatment for 48 hours (once each 24 hours) at a final concentration of 1ug/ml. Compounds were added to the plates to final concentration of 1uM for one hour and then cells were harvested by scraping. The samples were washed 2X in ice cold PBS, lysed (20mM Hepes, pH 7.5, 1% NP40, 1X HALT protease inhibitor cocktail, 1:1000 Pierce universal nuclease for cell lysis) on ice for 60 minutes with occasional mixing and centrifuged at 14k g for 15min to pellet cell debris. A 20ul aliquot was removed from each tube for 660nm protein assay for normalization.

2.7ml of Streptavidin Mag Sepharose beads (Cytiva# 28985799) were washed three times with 10ml, resuspended in 2.7ml lysis buffer and aliquoted between the nine samples equally. Samples were rotated end over end at room temperature for two hours. Following the binding step, beads were collected on the magnet, washed three times with lysis buffer, one time with water and eluted in 100ul of Hexafluoroisopropanol (Sigma #105228) for 5 minutes shaking at room temperature. Finally, eluted proteins were dried out for 10 minutes in speedvac, resuspended in 20ul of 1X LDS sample buffer with 1X NuPAGE™ sample reducing agent and boiled at 95°C for 5 minutes. Boiled samples were separated on a gel, transferred to a nitrocellulose mini membrane, and blocked with blocking buffer for 30 minutes.

Anti-GSK3 alpha/beta Monoclonal Antibody (Invitrogen #44-610) was added to the blot 1:1000 in PBS buffer with 0.2% Tween 20 (Sigma#P9416) to the blot for an hour shacking at room temperature. Blot wash washed four times (each 5 minutes) with PBS followed by one hour incubation with 1:4000 HRP conjugated mouse secondary antibody for an hour and four times PBS washes. Finally, the blot was treated with SuperSignal™ West Pico PLUS Chemiluminescent Substrate (ThermoFisher Scientific #34580) for 5 minutes and analyzed by BioRad ChemidDoc MP imaging system.

**Magnetic Bead Washing and Trypsin Digestion**

Beads were washed once with 1mL water, reduced with Reducing Buffer (10mM DTT, 100mM Tris, 1% SDS, pH8) at 70°C, and alkylated with 40mM iodoacetamide in Wash Buffer 1 (100mM Tris, 1% SDS, pH8) using a Kingfisher Duo Prime. Beads were further washed 3 times with 1mL Wash Buffer 1 (100mM Tris, 1% SDS, pH8), 5X cycles of 1mL Wash Buffer 2 (100mM Tris, 8M Urea, pH 8), and 5X cycles of 1mL Wash Buffer 3 (20% ACN) on a Kingfisher. Beads were transferred into 100uL Digestion Buffer (100mM TEAB, pH8, 10% ACN), and the proteins were digested with trypsin (1ug) overnight at 37°C on a Thermomixer while shaking.

**TMT Labeling, and high pH Fractionation**

Peptides were labeled with TMT 10plex reagent for 1hr, quenched with hydroxylamine, and pooled together. The solution was dried in a speedvac, then re-suspended in 100uL Buffer A (10mM ammonium Hydroxide, pH10), and subsequently injected onto a high pH fractionation system with a fraction collector using a Waters XBridge BEH130 C18 5um 2.1x100 mm column to separate the peptide mixture. A 60min gradient from 3-40%B (10mM ammonium Hydroxide in 90% ACN) was employed and 1min fractions were collected.

**LC-MS/MS**

Fractions from high pH system were dried in a speedvac, re-suspended in 0.1% formic acid, and loaded onto Evotips using the manufacturer's instructions. An Evosep One interfaced to a Lumos mass spectrometer (ThermoFisher Scientific) was used for all analyses. The standard Evosep 30 SPD method was used as well as the EV1106 column with Pepsep sprayer and fused silica emitter. The default TMT MS2 acquisition method was used.

**HTRF**

To evaluate the linker size on trimeric complex formation, Homogeneous Time Resolved Fluorescence (HTRF) assay was done using HaloTag standard protein (Promega #G449A) and in-house produced biotinylated DYRK1a kinase domain protein. The HaloTag was bound to the target protein using four compounds with 21, 30, 36 and 48 linker atoms in 7pt dose response (5uM maximal concentration with 3X dilution in between concentrations). After an hour of complex formation, Anti GST-Tb cryptate (Cisbio# 61GSTTLA) as a donor and Streptavidin-XLent (Cisbio #611SAXLA) as an acceptor were added to the samples according to manufacturer recommendation and incubated for an hour at room temperature. The assay was then read on a PerkinElmer EnVision Multimode Plate Reader.

**Chloroalkane penetration assay**

Six 12 well plates were seeded with TurboID-Halo cells and treated with 1ug/mL doxycycline for 24hr to initiate construct expression. Compounds in 6pt dose response (100uM maximal concentration with 4X dilution in between concentrations) were then added and incubated for 2, 6, or 24hr. All wells were harvested at the same time, and cells were washed 1X with ice cold PBS. 40uL of RIPA Buffer (Pierce) was added and mixed occasionally on ice for 1hr. The lysate was centrifuged for 15min at 14k g to pellet debris, and the supernatant was transferred to a fresh Eppendorf tube, and a small aliquot was used for a 660nM protein assay. The protein amounts were normalized between samples, and TMR-Halo ligand was added (1:1000) and incubated for 1hr at 37°C. Samples were prepared and separated on SDS Page mini-gels (Invitrogen) according to manufacturer's instructions. A Chemidoc imager (Biorad) was used for in-gel fluorescence measurements, and resulting images were processed in imageJ, and intensities were exported to Excel. $CP_{50}$ values were determined in GraphPad.

**Kinase selectivity profiling using quantitative chemical proteomics:**

**Affinity matrix generation**

24 X 500uL vials of Cytiva NHS activated magnetic sepharose beads were washed with 5ml DMSO (Dimethylsulfoxide) and beads were equilibrated for 15min at room temperature in DMSO. Beads were split into 2X aliquots, and pelleted by centrifugation, and the liquid was removed. DMSO was added to 1.8mL mark of each tube, and 36uL of 50mM compound (**1**)[1] was added to one batch of beads, and 36uL of 50mM of compound (**2**)[2] was added to the other batch of beads. 15 µl of triethylamine (SIGMA, T-0886, 99% pure) was added to each reaction to initiate conjugation. Beads were incubated at room temperature in darkness on an end-over-end shaker for overnight. Non-reacted NHS-groups were blocked by addition of 45uL ethanolamine and reacted at room temperature on the end-over-end shaker over-night. The beads were washed with 5mL volume of isopropanol 3X and stored in isopropanol at -20C.

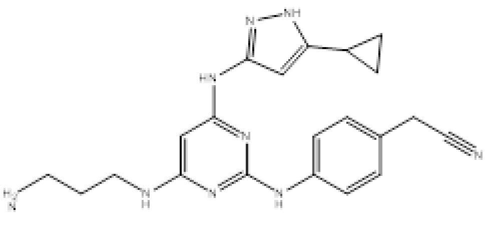
(**1**)

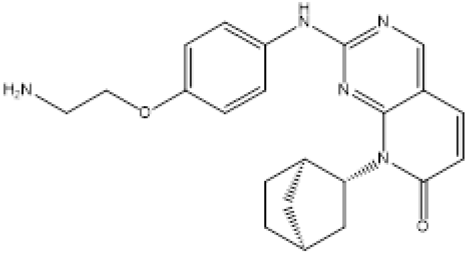
(**2**)

(1) Carrie M. Gower, Jason R. Thomas, Edmund Harrington, Jason Murphy, Matthew E. K. Chang, Ivan Cornella-Taracido, Rishi K. Jain, Markus Schirle, and Dustin J. Maly. Conversion of a Single Polypharmacological Agent into Selective Bivalent Inhibitors of Intracellular Kinase Activity. ACS Chemical Biology 2016 11 (1), 121-131. DOI: 10.1021/acschembio.5b00847
(2) Daub H, Olsen JV, Bairlein M, Gnad F, Oppermann FS, Körner R, Greff Z, Kéri G, Stemmann O, Mann M. Kinase-selective enrichment enables quantitative phosphoproteomics of the kinome across the cell cycle. Mol Cell. 2008 Aug 8;31(3):438-48. doi: 10.1016/j.molcel.2008.07.007. PMID: 18691976.

**HEK293 Lysate generation**

HEK293 cell pellets were thawed on ice and resuspended in 2X pellet volumes of lysis buffer (50mM Tris-HCL, 0.8% NP40, 5% glycerol, 150mM NaCl, 1.5mM MgCL2, 1mM DTT, pH7.5) supplemented with a protease inhibitor tablet (Complete EDTA-free, SIgma), and a phosphatase inhibitor tablet (Pierce, A32957). The sample was then homogenized in a dounce homogenizer, rotated for 30min at 4°C, centrifuged at 20,000g for 10min at 4°C, supernatant was removed, and aliquoted into 2 mL Eppendorf tubes for further use. Protein concentration was determined by a Pierce 660nM assay and aliquots were snap frozen at -80°C.

**Affinity enrichment and compound competition**

For each affinity enrichment condition, 500uL of HEK293 cell lysate (2.5 mg) was preincubated with varying concentrations (0.2nM-50uM) of $K_{PAL}$ for 1hr at 4°C in a 96 well Kingfisher deep well plate. During this preincubation, the derivatized magnetic sepharose beads (25 µL per sample) were washed 3X with Wash Buffer 2 (50 mM Tris pH 7.5, 150 mM NaCl, 1.5 mM MgCl2). Preincubated lysates were then incubated with derivatized resin for 1hr at 4°C with end-over-end agitation. A Kingfisher Duo Prime was used to wash beads with 1mL Wash Buffer 1 (50 mM Tris pH 7.5, 150 mM NaCl, 1.5 mM MgCl2, 0.4% IGEPAL CA-630) 3X, and then 1 mL Wash Buffer 2 2X. To elute bound proteins, beads were transferred into 50µL Elution buffer (10 mM DTT in 8M Urea/50mM TEAB) and incubated at 55°C for 30min. Proteins were alkylated with iodoacetamide (20mM) for 30min and diluted with 50mM TEAB to dilute Urea to 2M. Samples were on-bead digested with 1ug trypsin overnight.

**Mass spectrometry data acquisition and analysis**

Peptides were desalted using a uHLB SPE plate (Waters, MSPP-186001828) following the manufacturer's directions. After elution, samples were dried in a speedvac, and re-suspended in 50uL 50mM TEAB buffer for TMT labeling (Thermo Fisher) following the manufacturer's instructions. 20uL of TMT10plex reagents were added to each sample to encode the varying concentration of compound samples as well as the DMSO control. After quenching with hydroxylamine, the samples were combined and separated on an off-line high pH chromatography system (Agilent 1200 HPLC, Waters Xbridge column (2.1x100 mm) equipped with a fraction collector. The mobile phases and gradient consisted of A: 100% $H_2O$; B: 10mM $NH_4OH$; flow rate 0.2ml/min; 60min linear gradient from 1% mobile phase B to 40% mobile phase B. 1min fractions were collected in a 96 well plate and dried down in a speedvac. Fractions were re-suspended in 20uL 0.1% formic acid and loaded onto Evotips (Evosep). 60X 1min fractions were analyzed on an Evosep One connected to a Thermo Fisher Scientific Fusion Lumos mass spectrometer using the 30 sample per day method, and default TMT MS2 data acquisition method. Data was searched against a Uniprot human database using Proteome Discoverer (v2.5, Thermo Fisher Scientific) using 10ppm mass accuracy for the precursor mass, and 0.02 Da mass accuracy for the fragment ions, allowing for 2 missed tryptic cleavages. Carbamidomethyl (C) was selected as fixed modification and TMT6 (K), TMT6 (N-term), Oxidation (M) as variable modifications. GraphPad Prism was used to fit the 9-point dose response curves using non-linear regression.

**Chemical Synthesis of PAL and LMPL Probes:**

**$K_{PAL}$ Example 1:**

**3-(3-(but-3-yn-1-yl)-3H-diazirin-3-yl)-N-(4-(1-(tetrahydro-2H-pyran-4-yl)-1H-pyrrolo[2,3-c]pyridin-3-yl)pyridin-2-yl)propanamide (X)**

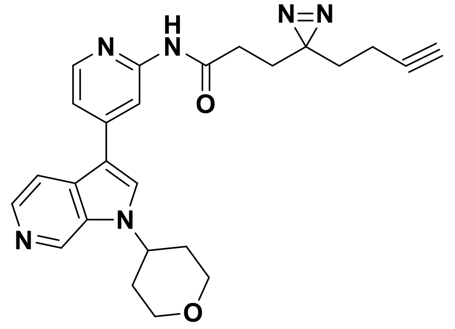

3-(3-(but-3-yn-1-yl)-3H-diazirin-3-yl)propanoic acid (18.0 mg, 0.108 mmol) was dissolved DCM (2 mL) and DMF (3 drops). To this was added oxalyl chloride (0.047 mL, 0.542 mmol). The mixture was stirred at RT for 2hr before concentrating and drying under vacuum for 30min. The residue was redissolved in DCM (2mL) and 4-(1-(tetrahydro-2H-pyran-4-yl)-1H-pyrrolo[2,3-c]pyridin-3-yl)pyridin-2-amine in DMF (2mL) was added, followed by Et3N (0.151mL, 1.08mmol) and stirred at RT for 2hr. The mixture was concentrated and purified by column chromatography with silica gel (eluent: 0-10% MeOH/DCM) to give a mixture that was further purified by HPLC (10-35% ACN/water with TFA) after concentrating to give the product (15mg 23.6% yield).

**$K_1$ Example 1:**

**21-chloro-N-(4-(1-(tetrahydro-2H-pyran-4-yl)-1H-pyrrolo[2,3-c]pyridin-3-yl)pyridin-2-yl)-3,6,9,12,15-pentaoxahenicosanamide (X)**

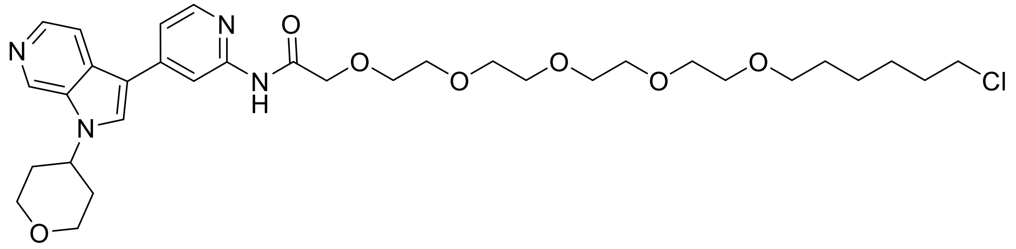

<u>Step 1:</u> 21-chloro-3,6,9,12,15-pentaoxahenicosanoic acid: A stirred solution of 21-chloro-3,6,9,12,15-pentaoxahenicosan-1-ol (200mg, 0.560mmol) in acetone (2.8mL) at RT was treated dropwise with Jones reagent (187µl, 1.12mmol). After stirring for 1hr at RT, MeOH (1mL) was added to the reaction mixture then diluted with EtOAc. The reaction mixture was washed with water, followed by brine, dried ($Na_2SO_4$), filtered, and concentrated. The resulting residue (190mg) was used without purification in the next step.

<u>Step 2:</u> 21-chloro-N-(4-(1-(tetrahydro-2H-pyran-4-yl)-1H-pyrrolo[2,3-c]pyridin-3-yl)pyridin-2-yl)-3,6,9,12,15-pentaoxahenicosanamide: A solution of 4-(1-(tetrahydro-2H-pyran-4-yl)-1H-pyrrolo[2,3-c]pyridin-3-yl)pyridin-2-amine (14.29mg, 0.049mmol), 21-chloro-3,6,9,12,15-pentaoxahenicosanoic acid (15mg, 0.040mmol), and HATU (21.53mg, 0.057mmol) in DMF (0.5mL) was treated with triethylamine (0.017mL, 0.121mmol) and stirred at RT for 16h. The reaction mixture was diluted with EtOAc and water, washed with brine, dried ($Na_2SO_4$), filtered, and concentrated. The resulting residue was purified by column chromatography over silica gel (4g, 0-15% MeOH/DCM) to afford the desired crude product which was re-purified by column chromatography (4g, 0-100% 3:1 EtOAc:EtOH/heptanes) to afford 21-chloro-N-(4-(1-(tetrahydro-2H-pyran-4-yl)-1H-pyrrolo[2,3-c]pyridin-3-yl)pyridin-2-yl)-3,6,9,12,15-pentaoxahenicosanamide (1.8mg, 6.53% yield).

**$K_2$ Example 1:**

**21-chloro-3,6,9,12,15-pentaoxahenicosyl 2-(2-(2-oxo-2-((4-(1-(tetrahydro-2H-pyran-4-yl)-1H-pyrrolo[2,3-c]pyridin-3-yl)pyridin-2-yl)amino)ethoxy)ethoxy)acetate**

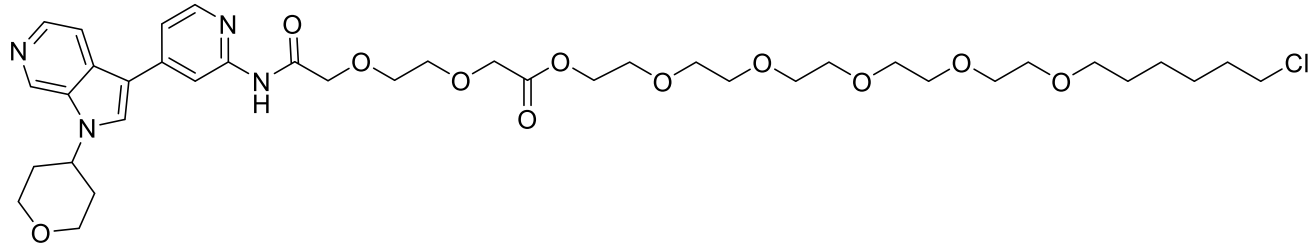

Step 1: 2-(2-(2-oxo-2-((4-(1-(tetrahydro-2H-pyran-4-yl)-1H-pyrrolo[2,3-c]pyridin-3-yl)pyridin-2-yl)amino)ethoxy)ethoxy)acetic acid: A solution of 4-(1-(tetrahydro-2H-pyran-4-yl)-1H-pyrrolo[2,3-c]pyridin-3-yl)pyridin-2-amine (20.0mg, 0.068mmol), 2,2'-(ethane-1,2-diylbis(oxy))diacetic acid (18.16mg, 0.102mmol), and HATU (51.7mg, 0.136mmol) in DMF (Volume: 0.5mL) was treated with triethylamine (0.028mL, 0.204mmol) and stirred at room temperature for 2 h. LCMS showed product formation. The reaction mixture was concentrated under a stream of nitrogen, was taken up in DCM:MeOH and was purified by silica gel chromatography (0-80% 3:1 EtOAc:EtOH / DCM; then 0-50% MeOH:DCM) to afford 2-(2-(2-oxo-2-((4-(1-(tetrahydro-2H-pyran-4-yl)-1H-pyrrolo[2,3-c]pyridin-3-yl)pyridin-2-yl)amino)ethoxy)ethoxy)acetic acid (8.8mg, 27% yield).

Step 2: 21-chloro-3,6,9,12,15-pentaoxahenicosyl 2-(2-(2-oxo-2-((4-(1-(tetrahydro-2H-pyran-4-yl)-1H-pyrrolo[2,3-c]pyridin-3-yl)pyridin-2-yl)amino)ethoxy)ethoxy)acetate: A solution of 2-(2-(2-oxo-2-((4-(1-(tetrahydro-2H-pyran-4-yl)-1H-pyrrolo[2,3-c]pyridin-3-yl)pyridin-2-yl)amino)ethoxy)ethoxy)acetic acid (8.8mg, 0.019mmol), 21-chloro-3,6,9,12,15-pentaoxahenicosan-1-ol (10.4mg, 0.029mmol), and HATU (14.7mg, 0.039mmol) in DMF (1mL) was treated with triethylamine (8.10µl, 0.058mmol) and stirred at room temperature for 16h. The reaction mixture was purified directly by HPLC (no TFA modifier) to afford 21-chloro-3,6,9,12,15-pentaoxahenicosyl 2-(2-(2-oxo-2-((4-(1-(tetrahydro-2H-pyran-4-yl)-1H-pyrrolo[2,3-c]pyridin-3-yl)pyridin-2-yl)amino)ethoxy)ethoxy)acetate as the product (5.0mg, 30.9% yield).

**$K_3$ Example 1:**

**21-chloro-N-(14-oxo-14-((4-(1-(tetrahydro-2H-pyran-4-yl)-1H-pyrrolo[2,3-c]pyridin-3-yl)pyridin-2-yl)amino)-3,6,9,12-tetraoxatetradecyl)-3,6,9,12,15-pentaoxahenicosanamide**

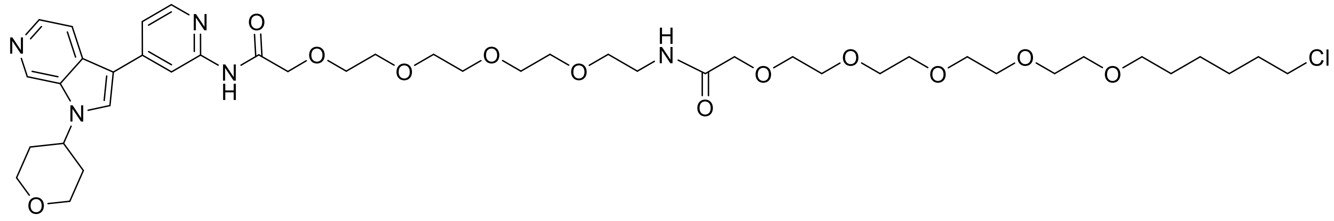

Step 1: (9H-fluoren-9-yl)methyl (14-oxo-14-((4-(1-(tetrahydro-2H-pyran-4-yl)-1H-pyrrolo[2,3-c]pyridin-3-yl)pyridin-2-yl)amino)-3,6,9,12-tetraoxatetradecyl)carbamate, 14-amino-N-(4-(1-(tetrahydro-2H-pyran-4-yl)-1H-pyrrolo[2,3-c]pyridin-3-yl)pyridin-2-yl)-3,6,9,12-tetraoxatetradecanamide: To a solution of 4-(1-(tetrahydro-2H-pyran-4-yl)-1H-pyrrolo[2,3-c]pyridin-3-yl)pyridin-2-amine (20mg, 0.068mmol), 1-(9H-fluoren-9-yl)-3-oxo-2,7,10,13,16-pentaoxa-4-azaoctadecan-18-oic acid (48.3mg, 0.102mmol), and HATU (51.7mg, 0.136mmol) in DMF (Volume: 0.5mL) was added triethylamine (0.028mL, 0.204mmol) and was stirred at room temperature for 16h. The reaction mixture was treated with piperidine (3 drops) was stirred at room temperature for 30 minutes. Additional piperidine (3 drops) was added and the reaction was stirred at room temperature for 1h, LCMS indicated the reaction was complete. The reaction mixture was concentrated under a stream of nitrogen and the resulting residue was taken up in DCM:MeOH and purified by column chromatography with silica gel (0-100% MeOH/DCM) to afford 14-amino-N-(4-(1-(tetrahydro-2H-pyran-4-yl)-1H-pyrrolo[2,3-c]pyridin-3-yl)pyridin-2-yl)-3,6,9,12-tetraoxatetradecanamide (48.0mg, 0.091mmol) and used without further purification.

Step 2: 21-chloro-N-(14-oxo-14-((4-(1-(tetrahydro-2H-pyran-4-yl)-1H-pyrrolo[2,3-c]pyridin-3-yl)pyridin-2-yl)amino)-3,6,9,12-tetraoxatetradecyl)-3,6,9,12,15-pentaoxahenicosanamide: To a solution of 14-amino-N-(4-(1-(tetrahydro-2H-pyran-4-yl)-1H-pyrrolo[2,3-c]pyridin-3-yl)pyridin-2-yl)-3,6,9,12-tetraoxatetradecanamide (48.0mg, 0.091mmol), 21-chloro-3,6,9,12,15-pentaoxahenicosanoic acid (67.5mg, 0.182mmol), and HATU (104mg, 0.273mmol) in DMF (1 mL) was added triethylamine (0.063mL, 0.455mmol) and the mixture was stirred at room temperature for 16h. The reaction mixture was purified directly by HPLC (no TFA modifier) to afford 21-chloro-N-(14-oxo-14-((4-(1-(tetrahydro-2H-pyran-4-yl)-1H-pyrrolo[2,3-c]pyridin-3-yl)pyridin-2-yl)amino)-3,6,9,12-tetraoxatetradecyl)-3,6,9,12,15-pentaoxahenicosanamide as the product (24.3mg, 28.8% yield).

**$K_4$ Example 1:**

**26-(21-chloro-3,6,9,12,15-pentaoxahenicosanamido)-N-(4-(1-(tetrahydro-2H-pyran-4-yl)-1H-pyrrolo[2,3-c]pyridin-3-yl)pyridin-2-yl)-3,6,9,12,15,18,21,24-octaoxahexacosanamide**

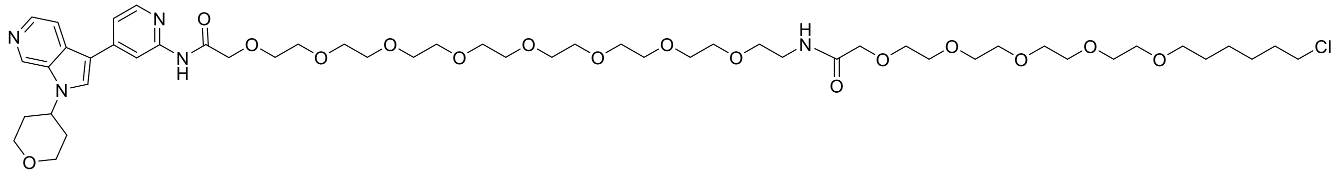

<ins>Step 1:</ins> (9H-fluoren-9-yl)methyl (26-oxo-26-((4-(1-(tetrahydro-2H-pyran-4-yl)-1H-pyrrolo[2,3-c]pyridin-3-yl)pyridin-2-yl)amino)-3,6,9,12,15,18,21,24-octaoxahexacosyl)carbamate, 26-amino-N-(4-(1-(tetrahydro-2H-pyran-4-yl)-1H-pyrrolo[2,3-c]pyridin-3-yl)pyridin-2-yl)-3,6,9,12,15,18,21,24-octaoxahexacosanamide

To a solution of 4-(1-(tetrahydro-2H-pyran-4-yl)-1H-pyrrolo[2,3-c]pyridin-3-yl)pyridin-2-amine (20mg, 0.068mmol), 1-(9H-fluoren-9-yl)-3-oxo-2,7,10,13,16,19,22,25,28-nonaoxa-4-azatriacontan-30-oic acid (66.2mg, 0.102mmol), and HATU (51.7mg, 0.136mmol) in DMF (0.5mL) was The reaction mixture was concentrated under a stream of nitrogen and was taken up in DCM/MeOH and was purified by silica gel chromatography (0-80% 3:1 EtOAc:EtOH / DCM; then 10-50% MeOH / DCM) to afford 26-amino-N-(4-(1-(tetrahydro-2H-pyran-4-yl)-1H-pyrrolo[2,3-c]pyridin-3-yl)pyridin-2-yl)-3,6,9,12,15,18,21,24-octaoxahexacosanamide as the product (22mg, 35.0% yield) by LCMS.

To a solution of 26-amino-N-(4-(1-(tetrahydro-2H-pyran-4-yl)-1H-pyrrolo[2,3-c]pyridin-3-yl)pyridin-2-yl)-3,6,9,12,15,18,21,24-octaoxahexacosanamide (22mg, 0.024mmol) in 0.5mL DCM was added piperidine (8 drops) and then stirred at room temperature for 20h. The reaction was concentrated under a stream of nitrogen and the resulting residue was azetroped with toluene three times then dried under vacuum to afford 26-amino-N-(4-(1-(tetrahydro-2H-pyran-4-yl)-1H-pyrrolo[2,3-c]pyridin-3-yl)pyridin-2-yl)-3,6,9,12,15,18,21,24-octaoxahexacosanamide as the product (31mg, 64.8%).The crude material was used directly in the next step.

<ins>Step 2:</ins> 26-(21-chloro-3,6,9,12,15-pentaoxahenicosanamido)-N-(4-(1-(tetrahydro-2H-pyran-4-yl)-1H-pyrrolo[2,3-c]pyridin-3-yl)pyridin-2-yl)-3,6,9,12,15,18,21,24-octaoxahexacosanamide

To a solution of 26-amino-N-(4-(1-(tetrahydro-2H-pyran-4-yl)-1H-pyrrolo[2,3-c]pyridin-3-yl)pyridin-2-yl)-3,6,9,12,15,18,21,24-octaoxahexacosanamide (31.0mg, 0.044mmol), 21-chloro-3,6,9,12,15-pentaoxahenicosanoic acid (82.0mg, 0.220mmol), and HATU (100mg, 0.264mmol) in DMF (1mL) was added triethylamine (0.061mL, 0.440mmol) and then stirred at room temperature for 16h. The reaction mixture was purified directly by HPLC (no TFA modifier) to afford 26-(21-chloro-3,6,9,12,15-pentaoxahenicosanamido)-N-(4-(1-(tetrahydro-2H-pyran-4-yl)-1H-pyrrolo[2,3-c]pyridin-3-yl)pyridin-2-yl)-3,6,9,12,15,18,21,24-octaoxahexacosanamide as the product (8.5mg, 17.4% yield).

**$K_5$ Example 1:**
**N-(4-(((2-amino-9H-purin-6-yl)oxy)methyl)benzyl)-2-(2-(2-(2-oxo-2-((4-(1-(tetrahydro-2H-pyran-4-yl)-1H-pyrrolo[2,3-c]pyridin-3-yl)pyridin-2-yl)amino)ethoxy)ethoxy)ethoxy)acetamide**

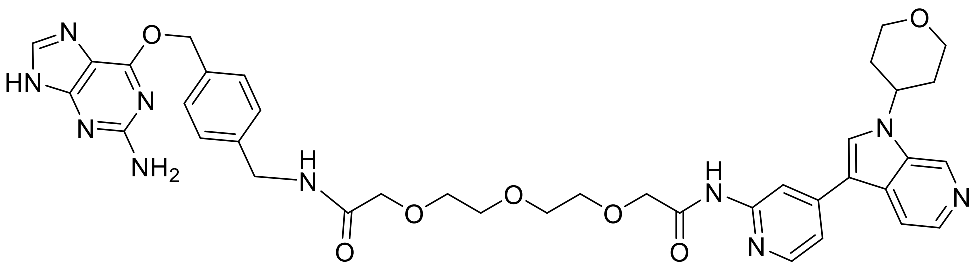

To a solution of 4-(1-(tetrahydro-2H-pyran-4-yl)-1H-pyrrolo[2,3-c]pyridin-3-yl)pyridin-2-amine (15.0 mg, 51.0 µmol), 1-(4-(((2-amino-9H-purin-6-yl)oxy)methyl)phenyl)-3-oxo-5,8,11-trioxa-2-azatridecan-13-oic acid (24.0mg, 51.0µmol), and HATU (19.0mg, 51.0µmol) in DMF (1.5mL) was added diisopropylethyl amine (6.6mg, 8.9µL, 51.0µmol) and stirred at room temperature for 1 h. Additional HATU (19.0mg, 51.0µmol) was added and the reaction mixture was stirred at room temperature for 20h. The reaction mixture was diluted with DMSO (1mL), filtered, then purified by HPLC. The product containing fractions were combined and lyophilized to afford N-(4-(((2-amino-9H-purin-6-yl)oxy)methyl)benzyl)-2-(2-(2-(2-oxo-2-((4-(1-(tetrahydro-2H-pyran-4-yl)-1H-pyrrolo[2,3-c]pyridin-3-yl)pyridin-2-yl)amino)ethoxy)ethoxy)ethoxy)acetamide as the trifluoroacetate salt (5.8mg, 13% yield).

**$K_6$ Example 1:**

**N-(4-(((2-amino-9H-purin-6-yl)oxy)methyl)benzyl)-14-(2-(2-(2-(2-oxo-2-((4-(1-(tetrahydro-2H-pyran-4-yl)-1H-pyrrolo[2,3-c]pyridin-3-yl)pyridin-2-yl)amino)ethoxy)ethoxy)ethoxy)acetamido)-3,6,9,12-tetraoxatetradecanamide**

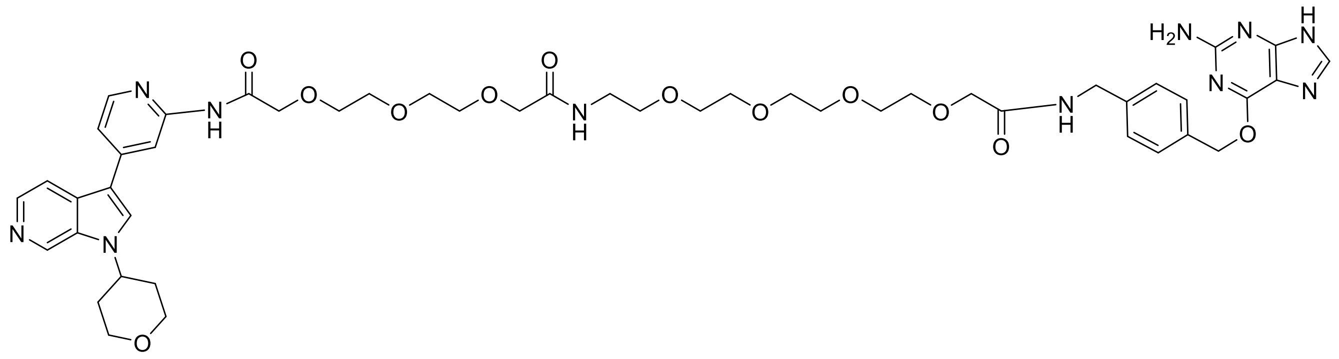

Step 1: (9H-fluoren-9-yl)methyl (1-(4-(((2-amino-9H-purin-6-yl)oxy)methyl)phenyl)-3-oxo-5,8,11,14-tetraoxa-2-azahexadecan-16-yl)carbamate: To a solution of 6-((4-(aminomethyl)benzyl)oxy)-9H-purin-2-amine (100mg, 1 Eq, 370µmol), 1-(9H-fluoren-9-yl)-3-oxo-2,7,10,13,16-pentaoxa-4-azaoctadecan-18-oic acid (175mg, 370µmol), and HATU (169mg, 1.2 Eq, 444µmol) in THF (10mL) was added diisopropylethyl amine (95.6mg, 129µL, 740µmol). The solution was then warmed to 50°C for 1h. The reaction mixture was cooled then concentrated under reduced pressure, diluted in DMSO and purified by reverse phase silica gel chromatography to afford (9H-fluoren-9-yl)methyl (1-(4-(((2-amino-9H-purin-6-yl)oxy)methyl)phenyl)-3-oxo-5,8,11,14-tetraoxa-2-azahexadecan-16-yl)carbamate (129mg, 48.0% yield) as the product.

Step 2: 14-amino-N-(4-(((2-amino-9H-purin-6-yl)oxy)methyl)benzyl)-3,6,9,12-tetraoxatetradecanamide, 1-(4-(((2-amino-9H-purin-6-yl)oxy)methyl)phenyl)-3,18-dioxo-5,8,11,14,20,23,26-heptaoxa-2,17-diazaoctacosan-28-oic acid: To a solution of (9H-fluoren-9-yl)methyl (1-(4-(((2-amino-9H-purin-6-yl)oxy)methyl)phenyl)-3-oxo-5,8,11,14-tetraoxa-2-azahexadecan-16-yl)carbamate (129mg, 178µmol) in DMF (5mL) was added DBU (135mg, 889µmol). The solution was then stirred at room temperature for 1h. The reaction mixture was treated with 2,2'-((oxybis(ethane-2,1-diyl))bis(oxy))diacetic acid (47.4mg, 213µmol) and HATU (81.1mg, 213µmol) and stirred at room temperature for 30 minutes. The reaction mixture was filtered and purified by HPLC to afford the product 1-(4-(((2-amino-9H-purin-6-yl)oxy)methyl)phenyl)-3,18-dioxo-5,8,11,14,20,23,26-heptaoxa-2,17-diazaoctacosan-28-oic acid as the trifluoroacetate salt (35mg, 24% yield).

Step 3: N-(4-(((2-amino-9H-purin-6-yl)oxy)methyl)benzyl)-14-(2-(2-(2-(2-oxo-2-((4-(1-(tetrahydro-2H-pyran-4-yl)-1H-pyrrolo[2,3-c]pyridin-3-yl)pyridin-2-yl)amino)ethoxy)ethoxy)ethoxy)acetamido)-3,6,9,12-tetraoxatetradecanamide: To a solution of 1-(4-(((2-amino-9H-purin-6-yl)oxy)methyl)phenyl)-3,18-dioxo-5,8,11,14,20,23,26-heptaoxa-2,17-diazaoctacosan-28-oic acid (4.7mg, 6.6μmol), 4-(1-(tetrahydro-2H-pyran-4-yl)-1H-pyrrolo[2,3-c]pyridin-3-yl)pyridin-2-amine (2.0mg, 6.6μmol), and HATU (3.0mg, 8.0μmol) in DMF (1mL) was added diisopropylethyl amine (2.6mg, 3.5μL, 20μmol) and was stirred at room temperature for 1h. HATU (3.0mg, 1.2 Eq, 8.0μmol) was added and the reaction was stirred at room temperature for 20h. Additional 4-(1-(tetrahydro-2H-pyran-4-yl)-1H-pyrrolo[2,3-c]pyridin-3-yl)pyridin-2-amine (2.0mg, 6.6μmol) was added and the reaction mixture was stirred at room temperature for 4 days. The reaction mixture was filtered and was purified by HPLC to afford N-(4-(((2-amino-9H-purin-6-yl)oxy)methyl)benzyl)-14-(2-(2-(2-(2-oxo-2-((4-(1-(tetrahydro-2H-pyran-4-yl)-1H-pyrrolo[2,3-c]pyridin-3-yl)pyridin-2-yl)amino)ethoxy)ethoxy)ethoxy)acetamido)-3,6,9,12-tetraoxatetradecanamide as the trifluioroacetate salt (0.6mg, 7% yield).

**$K_7$ Example 1:**

**N-(4-(((2-amino-9H-purin-6-yl)oxy)methyl)benzyl)-26-(2-(2-(2-(2-oxo-2-((4-(1-(tetrahydro-2H-pyran-4-yl)-1H-pyrrolo[2,3-c]pyridin-3-yl)pyridin-2-yl)amino)ethoxy)ethoxy)ethoxy)acetamido)-3,6,9,12,15,18,21,24-octaoxahexacosanamide**

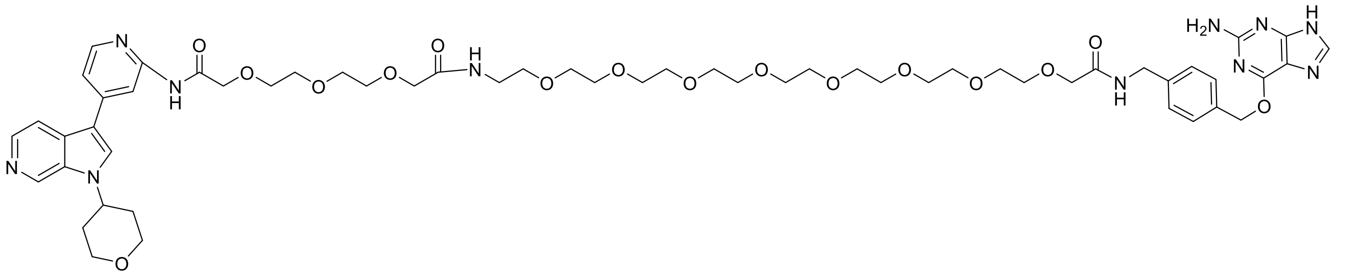

Step 1: (9H-fluoren-9-yl)methyl (1-(4-(((2-amino-9H-purin-6-yl)oxy)methyl)phenyl)-3-oxo-5,8,11,14,17,20,23,26-octaoxa-2-azaoctacosan-28-yl)carbamate: To a solution of 6-((4-(aminomethyl)benzyl)oxy)-9H-purin-2-amine (50mg, 0.18mmol), 1-(9H-fluoren-9-yl)-3-oxo-2,7,10,13,16,19,22,25,28-nonaoxa-4-azatriacontan-30-oic acid (0.12g, 0.18mmol), and HATU (84mg, 1.2 Eq, 0.22mmol) in DMF (2mL) was added diisopropylethylamine (64μL, 0.37mmol) and was stirred at room temperature for 3 days. The reaction mixture was concentrated under a stream of nitrogen, taken up in DMSO then purified by RP FCC to afford (9H-fluoren-9-yl)methyl

(1-(4-(((2-amino-9H-purin-6-yl)oxy)methyl)phenyl)-3-oxo-5,8,11,14,17,20,23,26-octaoxa-2-azaoctacosan-28-yl)carbamate (56mg, 34% yield) by LCMS.

Step 2: 26-amino-N-(4-(((2-amino-9H-purin-6-yl)oxy)methyl)benzyl)-3,6,9,12,15,18,21,24-octaoxahexacosanamide, 1-(4-(((2-amino-9H-purin-6-yl)oxy)methyl)phenyl)-3,30-dioxo-5,8,11,14,17,20,23,26,32,35,38-undecaoxa-2,29-diazatetracontan-40-oic acid: To a solution of (9H-fluoren-9-yl)methyl (1-(4-(((2-amino-9H-purin-6-yl)oxy)methyl)phenyl)-3-oxo-5,8,11,14,17,20,23,26-octaoxa-2-azaoctacosan-28-yl)carbamate (56.0mg, 62.0µmol) in DMF (2mL) was added DBU (47.0µL, 0.31mmol) and was stirred at room temperature. After 30 minutes 2,2'-((oxybis(ethane-2,1-diyl))bis(oxy))diacetic acid (28.0mg, 0.12mmol) and HATU (28.0mg, 74.0µmol) was added and the reaction mixture was stirred at RT for 20h. The reaction mixture was purified directly by RP FCC to afford 1-(4-(((2-amino-9H-purin-6-yl)oxy)methyl)phenyl)-3,30-dioxo-5,8,11,14,17,20,23,26,32,35,38-undecaoxa-2,29-diazatetracontan-40-oic acid (19.7mg, 36% yield) by LCMS.

Step 3: N-(4-(((2-amino-9H-purin-6-yl)oxy)methyl)benzyl)-26-(2-(2-(2-(2-oxo-2-((4-(1-(tetrahydro-2H-pyran-4-yl)-1H-pyrrolo[2,3-c]pyridin-3-yl)pyridin-2-yl)amino)ethoxy)ethoxy)ethoxy)acetamido)-3,6,9,12,15,18,21,24-octaoxahexacosanamide: To a solution of 1-(4-(((2-amino-9H-purin-6-yl)oxy)methyl)phenyl)-3,30-dioxo-5,8,11,14,17,20,23,26,32,35,38-undecaoxa-2,29-diazatetracontan-40-oic acid (6.0mg, 6.8µmol), 4-(1-(tetrahydro-2H-pyran-4-yl)-1H-pyrrolo[2,3-c]pyridin-3-yl)pyridin-2-amine (2.0mg, 6.8µmol), and HATU (3.1mg, 8.1µmol) in DMF (1mL) was treated with DIEA (2.6mg, 3.5µL, 3 Eq, 20µmol) and was stirred at RT for 20h. HATU (3.1mg, 8.1µmol) was added and the reaction was stirred at RT for an additional 24h. The reaction mixture was diluted with DMSO , filtered and was purified by HPLC to afford N-(4-(((2-amino-9H-purin-6-yl)oxy)methyl)benzyl)-26-(2-(2-(2-(2-oxo-2-((4-(1-(tetrahydro-2H-pyran-4-yl)-1H-pyrrolo[2,3-c]pyridin-3-yl)pyridin-2-yl)amino)ethoxy)ethoxy)ethoxy)acetamido)-3,6,9,12,15,18,21,24-octaoxahexacosanamide as the trifluoroacetate salt (1.5mg, 16% yield).

**$S_1$ Example 1:**

**(R)-1-(3-(4-((2-(2-((6-chlorohexyl)oxy)ethoxy)ethyl)amino)-4-oxobutanamido)phenyl)-3-(3,4-dimethoxyphenyl)propyl (S)-1-(3,3-dimethyl-2-oxopentanoyl)piperidine-2-carboxylate**

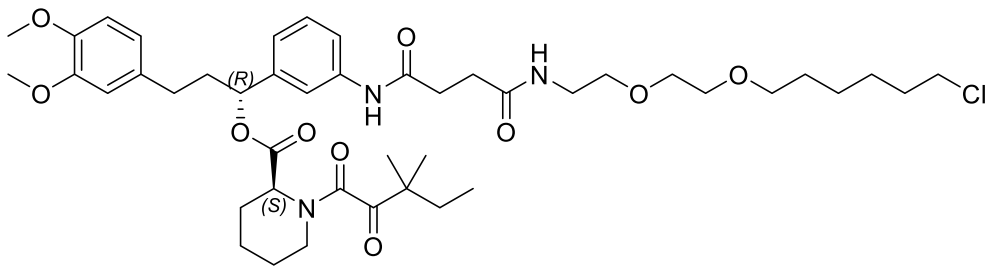

To a solution of Synthetic Ligand for FKBP (SLF) (15.0mg, 0.029mmol) in dry DMF (1mL) was added 4-((2-(2-((6-chlorohexyl)oxy)ethoxy)ethyl)amino)-4-oxobutanoic acid (10.2mg, 0.031mmol), HATU (13.05mg, 0.034mmol) and diisopropylethylamine (9.99µl, 0.057mmol) and the reaction mixture stirred at room temperature for 1h. The crude mixture was diluted with THF (1mL), filtered through PTFE membrane (0.20µm) and the resulting filtrate purified by HPLC to afford (R)-1-(3-(4-((2-(2-((6-chlorohexyl)oxy)ethoxy)ethyl)amino)-4-oxobutanamido)phenyl)-3-(3,4-dimethoxyphenyl)propyl (S)-1-(3,3-dimethyl-2-oxopentanoyl)piperidine-2-carboxylate as the product (14.8mg, 52.1%).

LC-MS (ES, *m/z*) = 830.6 [M+H]. $^1$H NMR (400 MHz, DMSO-d6): δ 9.97 (d, J = 5.3 Hz, 1H), 7.91 (t, J = 5.7 Hz, 1H), 7.67 (d, J = 17.9 Hz, 1H), 7.45 (d, J = 8.6 Hz, 1H), 7.28 (td, J = 7.9, 4.3 Hz, 1H), 7.01 (d, J = 7.5 Hz, 1H), 6.85 (d, J = 8.1 Hz, 1H), 6.80 - 6.74 (m, 1H), 6.68 (dd, J = 8.0, 2.0 Hz, 1H), 5.76 (s, 1H), 5.63 (dd, J = 8.7, 5.1 Hz, 1H), 5.14 (d, J = 5.5 Hz, 1H), 3.72 (d, J = 5.3 Hz, 7H), 3.61 (t, J = 6.6 Hz, 2H), 3.52 - 3.42 (m, 5H), 3.40 (d, J = 6.0 Hz, 2H), 3.29 (d, J = 3.6 Hz, 1H), 3.18 (dt, J = 9.8, 4.8 Hz, 3H), 2.39 (t, J = 7.1 Hz, 2H), 2.22 (d, J = 13.9 Hz, 1H), 2.01 (s, 1H), 1.63 (dtd, J = 32.7, 15.6, 14.6, 7.4 Hz, 8H), 1.48 (p, J = 6.8 Hz, 3H), 1.42 - 1.20 (m, 7H), 1.15 (d, J = 9.5 Hz, 6H), 1.04 (d, J = 4.2 Hz, 1H), 0.80 (t, J = 7.4 Hz, 3H):

**$S_2$ Example 1:**

**(R)-1-(3-(21-chloro-3,6,9,12,15-pentaoxahenicosanamido)phenyl)-3-(3,4-dimethoxyphenyl)propyl (S)-1-(3,3-dimethyl-2-oxopentanoyl)piperidine-2-carboxylate**

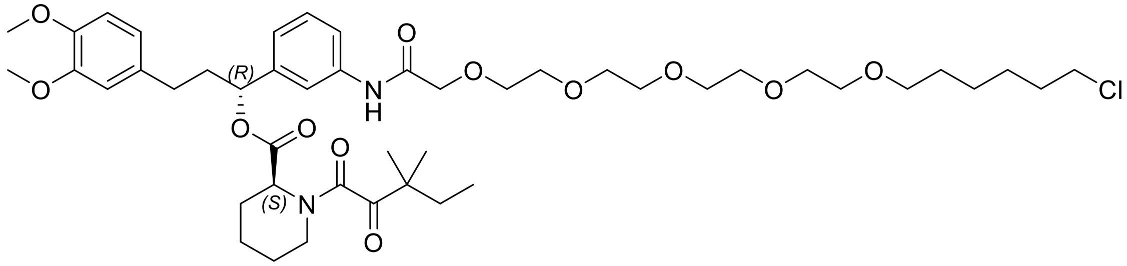

To a solution of SLF (20.0mg, 0.038mmol) in dry DMF (2mL) were added 21-chloro-3,6,9,12,15-pentaoxahenicosanoic acid (15.6mg, 0.042mmol) , HATU (17.39mg, 0.046mmol) and DIPEA (0.013mL, 0.076mmol). The reaction mixture was stirred at room temperature for 90 minutes. The crude mixture was filtered through PTFE membrane (0.20µm) and the resulting filtrate purified HPLC to afford for (R)-1-(3-(21-chloro-3,6,9,12,15-pentaoxahenicosanamido)phenyl)-3-(3,4-dimethoxyphenyl)propyl (S)-1-(3,3-dimethyl-2-oxopentanoyl)piperidine-2-carboxylate (14.6mg, 99% yield) as a yellow oil.

LC-MS (ES, *m/z*) = 894.3 [M+H]. 1H NMR (400 MHz, DMSO-d6): δ 9.65 (s, 1H), 7.77 - 7.67 (m, 1H), 7.60 - 7.50 (m, 1H), 7.31 (td, J = 7.9, 4.4 Hz, 1H), 7.06 (d, J = 7.5 Hz, 1H), 6.85 (d, J = 8.2 Hz, 1H), 6.77 (dd, J = 5.7, 2.0 Hz,
1H), 6.68 (dd, J = 8.2, 1.9 Hz, 1H), 5.66 (dd, J = 8.6, 5.0 Hz, 1H), 5.15 (d, J = 5.4 Hz, 1H), 4.26 (dd, J = 19.8, 9.1 Hz, 0H), 4.07 (s, 2H), 3.71 (d, J = 5.8 Hz, 6H), 3.66 (dd, J = 6.1, 3.3 Hz, 2H),3.63 - 3.51 (m, 8H), 3.47 (s, 6H), 3.38 - 3.27 (m, 3H), 2.54 - 2.47 (m, 8H), 2.47 (s, 8H), 1.74 - 1.58 (m, 6H), 1.50 (ddt, J = 31.8, 13.8, 6.9 Hz, 3H), 1.41 - 1.17 (m, 6H), 1.15 (d, J = 9.4 Hz, 5H), 1.04 (d, J = 4.3 Hz, 1H), 0.80 (t, J = 7.4 Hz, 2H).

**$S_3$ Example 1:**
**(R)-1-(3-(36-chloro-16-oxo-3,6,9,12,18,21,24,27,30-nonaoxa-15-azahexatriacontanamido)phenyl)-3-(3,4-dimethoxyphenyl)propyl (S)-1-(3,3-dimethyl-2-oxopentanoyl)piperidine-2-carboxylate**

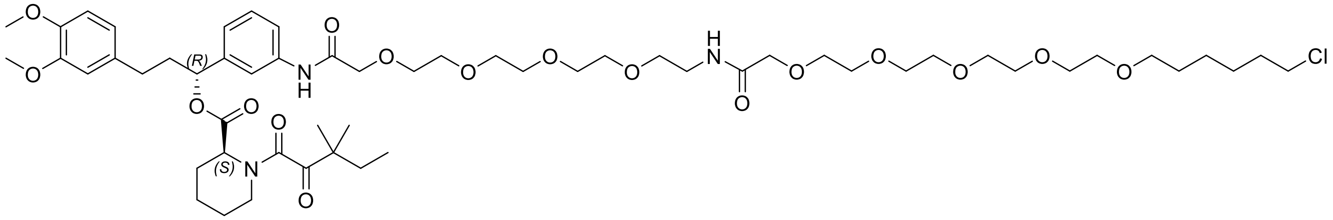

<u>Step 1:</u> (R)-1-(3-(1-(9H-fluoren-9-yl)-3-oxo-2,7,10,13,16-pentaoxa-4-azaoctadecan-18-amido)phenyl)-3-(3,4-dimethoxyphenyl)propyl (S)-1-(3,3-dimethyl-2-oxopentanoyl)piperidine-2-carboxylate: To a solution of SLF (8.9mg, 17.0µmol), 1-(9H-fluoren-9-yl)-3-oxo-2,7,10,13,16-pentaoxa-4-azaoctadecan-18-oic acid (9.6mg, 1.2 Eq, 20µmol), and HATU (9.7mg, 1.5 Eq, 25µmol) in DMF (0.5mL) at room temperature was added with diisopropylethylamine (4.4mg, 5.9µL, 34.0µmol) and the mixture stirred for 2h. The reaction mixture was purified directly by RP

FCC to afford (R)-1-(3-(1-(9H-fluoren-9-yl)-3-oxo-2,7,10,13,16-pentaoxa-4-azaoctadecan-18-amido)phenyl)-3-(3,4-dimethoxyphenyl)propyl (S)-1-(3,3-dimethyl-2-oxopentanoyl)piperidine-2-carboxylate as the product (15mg, 90% yield).

<u>Step 2:</u> (R)-1-(3-(36-chloro-16-oxo-3,6,9,12,18,21,24,27,30-nonaoxa-15-azahexatriacontanamido)phenyl)-3-(3,4-dimethoxyphenyl)propyl (S)-1-(3,3-dimethyl-2-oxopentanoyl)piperidine-2-carboxylate: To a solution of (R)-1-(3-(1-(9H-fluoren-9-yl)-3-oxo-2,7,10,13,16-pentaoxa-4-azaoctadecan-18-amido)phenyl)-3-(3,4-dimethoxyphenyl)propyl (S)-1-(3,3-dimethyl-2-oxopentanoyl)piperidine-2-carboxylate (15mg, 15µmol) in DCM (1mL) was treated with DBU (23µL, 0.15mmol) and stirred at room temperature for 1h. The reaction mixture was concentrated under a stream of nitrogen overnight. The resulting residue was taken up in DMF (0.5mL) and treated with 21-chloro-3,6,9,12,15-pentaoxahenicosanoic acid (8.5mg, 1.5 Eq, 23µmol), 2-(3H-[1,2,3]triazolo[4,5-b]pyridin-3-yl)-1,1,3,3-tetramethylisouronium hexafluorophosphate(V) (12.0 mg, 31.0 µmol) and diisopropylethylamine (13.0µL, 77.0µmol) then stirred for 3h. The reaction mixture was filtered and purified directly by to afford (R)-1-(3-(36-chloro-16-oxo-3,6,9,12,18,21,24,27,30-nonaoxa-15-azahexatriacontanamido)phenyl)-3-(3,4-dimethoxyphenyl)propyl (S)-1-(3,3-dimethyl-2-oxopentanoyl)piperidine-2-carboxylate, AI - Trifluoroacetate (4.0mg, 20% yield).
